# Targeting Lysosomal Dysfunction to Alleviate Plaque Deposition in an Alzheimer Disease Model

**DOI:** 10.1101/2025.04.28.651121

**Authors:** Leigh Ellen Fremuth, Diantha van de Vlekkert, Huimin Hu, Jason Andrew Weesner, Ida Annunziata, Gouri Yogalingam, Alessandra d’Azzo

## Abstract

Alzheimer disease (AD) is characterized by aberrant amyloid precursor protein (APP) processing and lysosomal dysfunction. This study identifies two members of the lysosomal multi-enzyme complex (LMC), neuraminidase 1 (Neu1) and protective protein/cathepsin A (PPCA), as a critical regulators of APP metabolism. Neu1 deficiency in human AD brains and 5xFAD/*Neu1^-/-^* mice leads to sialic acid retention on APP and its secretases, enhancing amyloidogenic cleavage and Aβ42 production. Additionally, Neu1 deficiency increases lysosomal exocytosis, contributing to extracellular Aβ release and neuroinflammation. Conversely, overexpression of PPCA in neurons or co-expression of PPCA and NEU1 normalizes sialylation patterns, reduces secretase activity, and mitigates plaque burden. These findings reveal a novel bidirectional dependency between Neu1 and PPCA, underscoring their cooperative role in maintaining lysosomal homeostasis. Additionally, AAV-mediated co-expression of NEU1 and PPCA in 5XFAD brains demonstrates therapeutic potential by reducing amyloid pathology. These findings position lysosomal dysfunction and the Neu1-PPCA axis as promising targets for therapeutic intervention in AD.

## Introduction

The amyloid precursor protein (APP) has been studied for over 50 years as a key factor in the pathology of amyloidosis, with most of the research focused on its role in Alzheimer disease (AD) pathogenesis. APP is a type-I transmembrane protein ubiquitously expressed across various tissues, particularly neuronal tissue. It is found in multiple subcellular compartments, including the plasma membrane (PM) and endo-lysosomal vesicles^1^. Studies investigating APP’s functions have uncovered its involvement in processes such as cell adhesion, neural-trophic functions, anterograde axonal transport, neural plasticity, synapse formation and repair, and iron export. However, the underlying mechanisms driving these functions remain only partially understood^1–5^.

In recent years, research has shifted from studying full-length APP to exploring the functional roles of and interactions of its proteolytic products, including soluble APP (sAPPs), APP C-terminal fragments (CTFs), amyloid β (Aβ) peptides, and APP intracellular domains (AICD). These fragments are generated by two distinct proteolytic processing pathways: the non-amyloidogenic and amyloidogenic cleavage of APP^6^. In the non-amyloidogenic pathway, APP is cleaved at the PM, first by α-secretases, generating sAPPα and α-CTF. α-CTF is then further processed by γ-secretase to yield P3 peptides and AICDs^6^. Functionally, sAPPα has been shown to promote neurogenesis and to exert neuroprotective effects, while AICDs have been implicated in apoptosis and transcriptional regulation. P3 peptides have been described as neuroprotective in some context but have also been shown to induce cytotoxicity through the formation of aggregates^1,7–9^. The biological functions of α-CTF, however, remain largely uncharacterized. In contrast, amyloidogenic processing of APP occurs within the endo-lysosomal system, where APP is initially cleaved by b-secretase into sAPPβ and β-CTF.

β-CTF is subsequently processed by γ-secretase to generate Aβ peptides and the same AICDs that are produced during non-amyloidogenic cleavage^6^. Functionally, sAPPβ have been shown to promote axon pruning and neurodegeneration, and recent studies have identified β-CTF as a contributor to synaptic toxicity, mitochondrial dysfunction, and lysosomal impairment. Nevertheless, most functional studies of amyloidogenic APP products have centered on Aβ peptides, which have been widely recognized as key pathogenic players in AD^1,10–12^. β-CTF can be cleaved at multiple sites by γ-secretase, generating a variety of Aβ peptides ranging from 37 to 49 amino acids in length, with Aβ40 and Aβ42 being the most extensively studied^12^. These peptides can be secreted into the extracellular space, where they aggregate into oligomers that mature into fibrils, leading to spine degeneration and synapse loss^6^. Although debate persists over the exact role of Aβ peptides in the development of AD, there is broad consensus that they contribute significantly to AD pathogenesis, particularly through their potent immunomodulatory effects^13,14^.

Therapeutic strategies for AD often focus on regulating the proteolytic processing of APP, aiming to modulate the levels of its cleavage products in the brain parenchyma^15^. Consequently, a detailed understanding of how APP processing is controlled at the enzymatic level is critical. Canonically, the primary α-secretase mediating the first step of non-amyloidogenic cleavage is tumor necrosis factor-alpha converting enzyme (TACE), whereas the main β-secretase driving amyloidogenic cleavage is beta-site APP-cleaving enzyme 1 (BACE1)^16^. The γ-secretase complex, which participates in both pathways, consists of four core proteins with distinct functions: presenilin (PS), the catalytic subunit; presenilin enhancer 2 (PEN2), which facilitates PS maturation; anterior pharynx-defective 1 (APH1), the scaffolding subunit; and nicastrin (NCT), reportedly required for γ-secretase substrate recognition and selection^16–18^.

Glycosylation, particularly sialylation, is a prominent post-translational modification that regulates enzyme stability, trafficking, and catalytic activity, yet it remains relatively understudied in the context of AD and the secretases^19^. The glycosylation process is among the most common and complex post-translational modifications, and sialylation - a specific type of glycosylation - refers to the addition of sialic acids, negatively charged sugars highly concentrated in the brain, to the terminal ends of glycan chains^19^. Aberrant glycosylation of APP and the secretases, as well as increased sialylation of microglial proteins, have been observed in the brains of AD patients, although these modifications have yet to be directly targeted therapeutically^20–22^. Altered glycosylation of APP has been shown to affect its transport and trafficking, while enhanced sialylation specifically has been linked to increase APP secretion and elevated Aβ production^21,23^. Similarly, glycosylation of BACE1 has been demonstrated to be required for its proper endosomal compartmentalization and enzymatic function^24,25^, and the glycosylation levels of TACE has also been shown to influence its enzymatic activity^26^. The role of glycosylation in γ-secretase activity remains a subject of debate, particularly regarding its substrate recognition subunit, NCT^27^. Some studies have reported that glycosylation of NCT does not impact the enzymatic function of γ-secretase; however, more recent findings suggest that NCT glycosylation is crucial for γ-secretase enzyme activity and substrate specificity^28,29^. Furthermore, alterations in the glycosylation of proteins that interact with or regulate the secretases can impact their biochemical properties and physiological functions^30^. Despite these insights, the broader impact of glycosylation on APP processing remains poorly understood, and studies specifically investigating the role of sialylation are exceedingly rare.

Aberrant glycosylation of APP and the secretases has been linked to lysosomal dysfunction, another prominent pathological feature in the progression of AD^30^. To date, most investigations have focused on alterations in the addition of glycans to APP and the secretases, a process that primarily occurs in the Golgi apparatus. However, another potential layer of regulation is the removal or modification of glycan residues (de-glycosylation), which typically takes place within the endo-lysosomal compartment and is mediated by lysosomal enzymes. Support for this additional regulatory mechanism comes from studies of lysosomal storage diseases (LSDs), a group of more than 70 disorders characterized by lysosomal dysfunction and pathology that closely mirrors what has been found to occur in major neurodegenerative diseases, including AD^31,32^.

While many groups have explored the parallels between adult neurogenerative diseases and LSDs, our group has specifically established similarities between the LSD sialidosis and AD. Sialidosis is a rare genetic disorder caused by mutations in the *neuraminidase 1* (*NEU1*) gene, which encodes the most ubiquitous sialidase responsible for cleaving terminal sialic acid residues from glycoproteins in the lysosome, and occasionally at the PM^33^. NEU1 operates as part of a multiprotein complex and depends on one of its components, the serine carboxypeptidase protective protein cathepsin A (PPCA), for proper lysosomal localization, enzyme stability and catalytic activity^34^. From our studies, we discovered that NEU1 functions as a negative regulator of lysosomal exocytosis, a process involving the docking and fusion of lysosomes with the PM, and the subsequent extracellular release of lysosomal contents, including active lysosomal enzymes^35^. In addition, through collaborative effort, we identified PPCA as a key regulator of chaperone-mediated autophagy (CMA)^36^. Both lysosomal exocytosis and CMA have since been recognized as pathogenic mechanisms in AD^37^. Building on these findings, we established that two major AD-related proteins - APP and Triggering Receptor Expressed on Myeloid Cells 2 (Trem2), a prominent microglial receptor - are substrates of NEU1. We demonstrated that insufficient de-sialylation of these proteins, due to *Neu1* deficiency or downregulation, drives AD-like pathology, including enhanced Aβ42 release, increased plaque formation, and neurotoxic, microglia-mediated inflammation^38,39^.

Given our previous findings, and the remaining gaps in understanding APP processing, we sought to investigate how Neu1-mediated de-sialylation contributes to the progression of amyloidosis pathology at the level of secretase function using 5xFAD mice, a well-established model of AD. Here, we identify two members of the lysosomal multi-enzyme complex, Neu1 and PPCA, as critical regulators of APP cleavage. Using AD patient tissue and 5xFAD mouse models, we show that Neu1 deficiency increases sialylation of APP and secretases, enhances lysosomal exocytosis, and accelerates amyloidogenic processing. Conversely, PPCA overexpression in neurons or NEU1/PPCA co-expression ameliorates sialylation levels, reduces secretase hyperactivity, and diminishes amyloid plaque burden, especially in female mice. Taken together, these data uncover a novel bidirectional dependency between Neu1 and PPCA, emphasizing that proper sialylation is essential for lysosomal and APP homeostasis. Overall, these findings highlight targeting lysosomal dysfunction in AD as a promising therapeutic intervention.

## Results

### NEU1 May Influence Amyloidogenic Enzymes in Alzheimer’s Disease Through Modulation of Their Sialylation

We began by assessing NEU1 expression levels in brain and hippocampus specimens from 20 AD patients, comparing them to 20 control samples. The total NEU1 protein levels were slightly reduced in the AD brain samples compared to the controls (Fig. 1a and Extended Data Fig. 1a). These findings aligned with data from microarray datasets^39^, further supporting the idea that the downregulation of NEU1 may play a significant role in AD pathology, even in the absence of a direct association in GWAS studies. One result of diminished NEU1 activity was the increased levels of sialylated PPCA protein observed in AD patients (Fig. 1b), with no corresponding increase in total protein levels (Extended Data Fig. S1b). These findings also indicate that PPCA is a substrate of NEU1, a correlation that has not yet been defined. A second indicator of reduced NEU1 levels was the extent of lysosomal exocytosis, which we measured by analyzing LAMP1 protein levels and extracellular β-Hexosaminidase (β-Hex) enzyme activity, a canonical lysosomal enzyme. In AD patients, we observed increased LAMP1 staining in hippocampal sections (Fig. 1c) along with elevated β-HEX activity in serum (Fig. 1d). These findings indicate that lysosomal exocytosis is enhanced in AD patients as a result of NEU1 downregulation. Further analysis of β-HEX activity in AD patients’ samples revealed that female patients were more affected (Fig. 1d). In addition, AD patients, but not controls, showed a significant increase in serum β-HEX activity with age (Fig. 1e), suggesting that a progressive rise in lysosomal exocytosis correlates with the worsening of AD pathology.

**Fig. 1:**
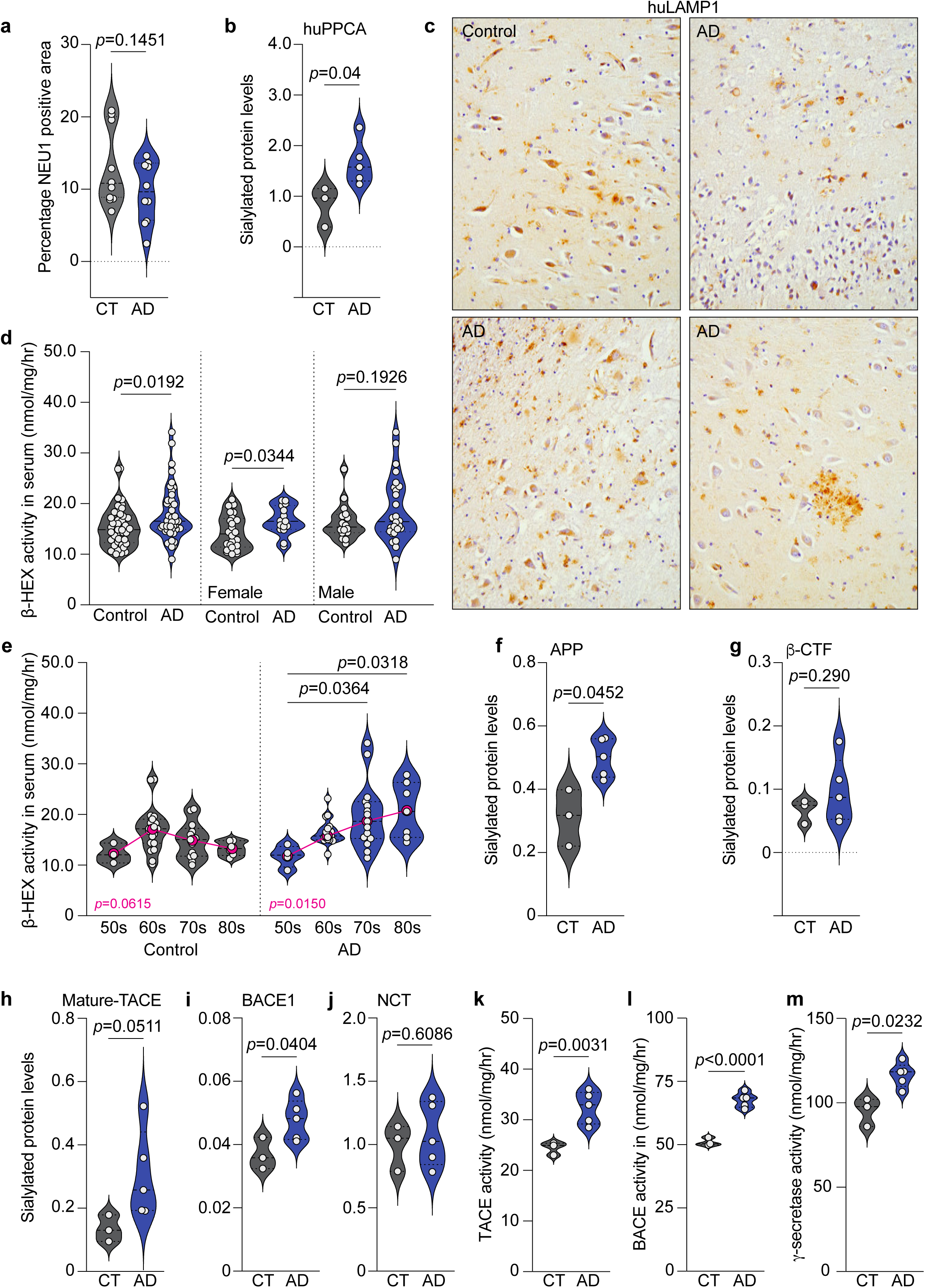
LMC- and AD-associated proteins in human AD patient material. **a**, Quantification of NEU1^+^ area per section area, displayed as a percentage, in human controls (n=10) and AD patients (n=10). **b**, Normalized expression of sialylated huPPCA, relative to total protein levels displayed in *Extended Data Fig. 1b*, in hippocampal tissue from human controls (n=3) and AD patients (n=5). **c**, Immunohistochemical staining of LAMP1 (brown) in human controls (n=10) and AD patients (n=10). **d**, Quantification of β-hexosaminidase (β-HEX) in serum collected from human controls (n=41) and AD (n=41) patients, plotted as all “n” (left), females only (n=24, n=15, middle), and males only (n=16, n=26, right). **e**, β-HEX quantification re-plotted by age as control 50s (n=3), control 60s (n=19), control 70s (n=13), control 80s (n=7), AD 50s (n=4), AD 60s (n=15), AD 70s (n=15) and AD 80s (n=7). **f-m**, Measurements collected from homogenized hippocampal tissue of controls (n=3) and AD patients (n=5). **f**, Normalized expression of sialylated APP, relative to total protein levels displayed in *Extended Data Fig. 1c*. **g**, Normalized expression of sialylated β-CTF, relative to total protein levels displayed in *Extended Data Fig. 1d*. **h**, Normalized expression of sialylated mature TACE, relative to total protein levels displayed in *Extended Data Fig. 1e*. **i**, Normalized expression of sialylated BACE1, relative to total protein levels displayed in *Extended Data Fig*. 1f. **j**, Normalized expression of sialylated NCT, relative to total protein levels displayed in *Extended Data Fig. 1g*. **k**, Quantification of TACE enzyme activity. **l**, Quantification of BACE1 enzyme activity. **m**, Quantification of γ-secretase enzyme activity. Actual p-values are given, any value less than or equal to 0.05 is considered significant.

Furthermore, we observed that APP behaved similarly to PPCA, with AD patients exhibiting higher levels of sialylated APP (Fig. 1f), despite total levels being comparable to those in controls (Extended Data Fig. 1c). These data are consistent with our previous findings, which showed that APP is a substrate of NEU1 and remains sialylated in the commonly used 5xFAD mouse model of AD^38^. In this study, we also observed the accumulation of the β-CTF fragment of APP^38^. We confirmed that this was also the case in AD patients, who exhibited higher levels of total β-CTF compared to controls (Extended data Fig. 1d). Previous studies have predicted that β-CTF could be glycosylated, carrying O-linked glycans rather than N-linked glycans^40,41^. We validated these suggestions by demonstrating that β-CTF is sialylated in both AD patients and controls, with AD patients showing a trend toward higher levels of sialylated protein, although this difference was not statistically significant (Fig. 1g).

It has been established that alterations in the glycosylation of APP will greatly impact its processing by secretases. Additionally, the glycosylation of these enzymes themselves has been implicated in modulating their activity and substrate processing. To investigate this further, we measured the total and sialylated levels of α-secretase (mature TACE), β-secretase (BACE1), and the substrate selection unit of γ-secretase, nicastrin (NCT). We found that while there were no significant differences in the total protein levels between AD patients and controls (Extended Data Figs. 1e-1g), there were increased levels in sialylated TACE and BACE1 (Figs. 1h and 1i). No differences were observed in the levels of sialylated NCT (Fig. 1j) and pro-TACE was undetectable in our assay. We next measured the enzyme activity of TACE, BACE1, and γ-secretase, all of which were significantly higher in AD patients compared to controls (Figs. 1k-1m). The increased levels of sialylated proteins, combined with the enhanced enzyme activities, imply that sialylation of APP, TACE, and BACE1 positively influences the enzyme activity of TACE and BACE, leading to enhanced processing of APP, while sialylation of β-CTF, but not NCT, may impact its processing by γ-secretase.

### Modeling Alzheimer’s Disease Pathology by Knocking Out Neu1 in 5xFAD Mice

In previous work, we showed that *Neu1*^-/-^ mice crossed with 5xFAD mice exhibited increased levels of sialylated APP in hippocampal tissue and elevated levels of amyloid peptides released into the CSF^38^. Building on these findings, which align with some of the human AD data presented in Fig. 1, we aimed to further characterize the Neu1 deficient/5xFAD mice (5xFAD/*Neu1*^-/-^) and explore additional parallels between this mouse model and human AD patients. Our goal was to establish the 5xFAD/*Neu1*^-/-^ mouse as an ideal model for studying lysosomal dysfunction in AD. We began by quantifying and characterizing the plaque burden in the brains of 5xFAD/ *Neu1*^-/-^ mice where Neu1 activity was significantly reduced (Extended Data Fig. 2a). A gender bias was immediately apparent: female 5xFAD/*Neu1*^-/-^ brains showed no difference in plaque burden compared to 5xFAD controls, while male brains exhibited a significant increase in plaque burden (Fig. 2a and Extended Data Fig. 2b). Interestingly, in the presence of normal Neu1 levels, female 5xFAD mice had a higher plaque burden than their male counterparts, a difference that was abolished when Neu1 was ablated (Fig. 2a). Next, we showed that Neu1 deficiency increased the average number of plaques per brain area in the 5xFAD brain, regardless of gender (Fig. 2b). However, it differentially affected average plague size: female plaque size tended to decrease, while male plaque size significantly increased (Fig. 2c). Notably, in the 5xFAD background, when Neu1 was present, there was no gender bias for plaque number, but female mice exhibited significantly larger average plaque sizes compared to males (Fig. 2b and 2c). One additional observation was that Neu1 deficiency, whether alone or in the 5xFAD background, resulted in a decrease in total brain size (Extended Data Fig. 2c). Overall, the addition of Neu1 deficiency to the 5xFAD background appears to differentially impact female and male mice, diminishing the gender biases observed in Neu1-sufficient 5xFAD cohorts.

**Fig. 2:**
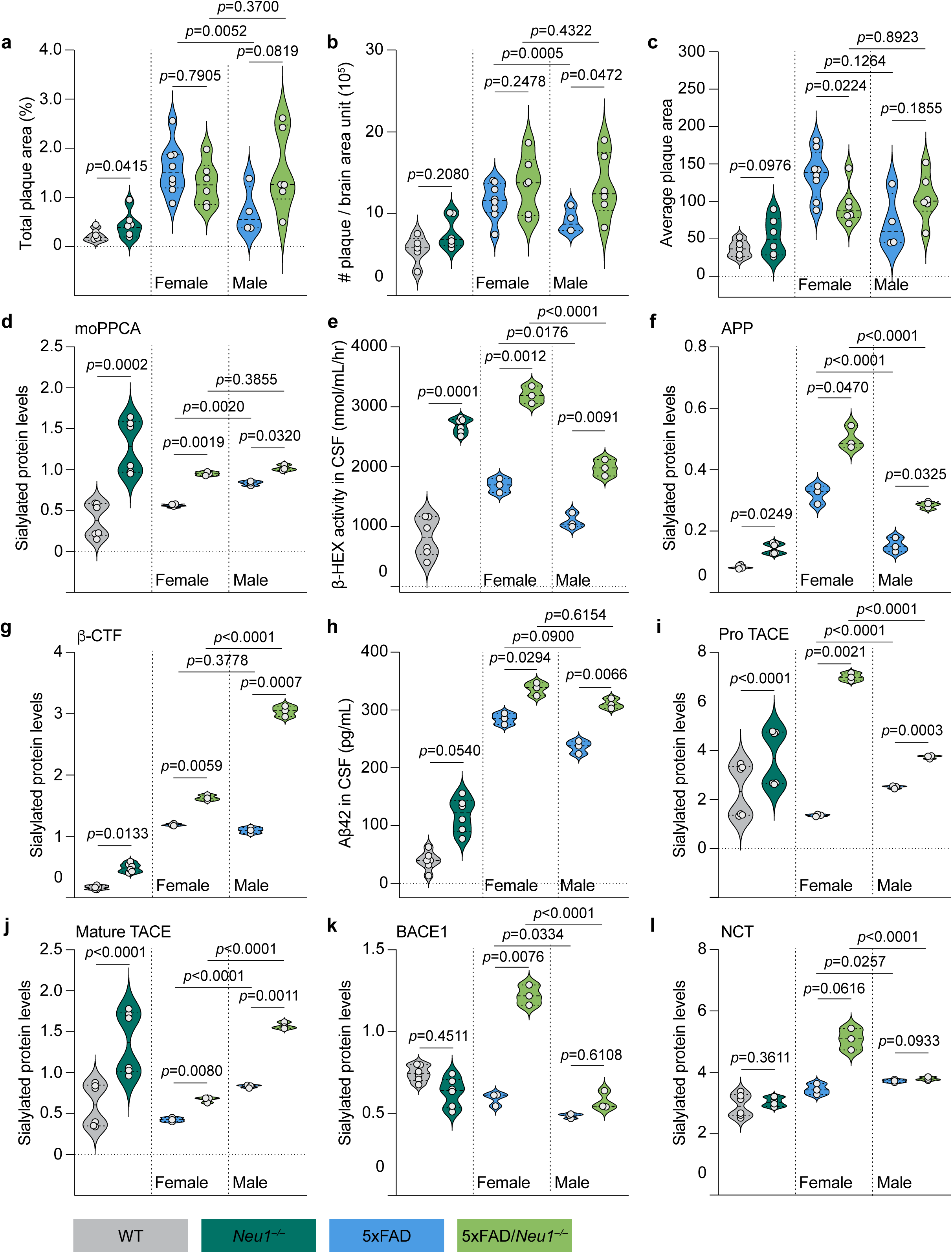
***Neu1* deficiency in 5xFAD mice. a-c**, Measurements collected from paraffin-embedded whole brain sections for WT (n=5), *Neu1*^-/-^ (n=6), female 5xFAD (n=8), female 5xFAD/*Neu1*^-/-^ (n=6), male 5xFAD (n=4) and male 5xFAD/*Neu1*^-/-^ (n=6) mice. **a**, Quantification of plaque burden, represented as amyloid^+^ (4G8^+^) area per section area, displayed as a percentage. **b**, Quantification of the average number of plaques per brain area unit (total number of plaques/total brain area). **c**, Quantification of the average plaque area (total plaque area/total plaque number). **d-l**, Measurements collected from WT (n=6), *Neu1*^-/-^ (n=6), female 5xFAD (n=3), female 5xFAD/*Neu1*^-/-^ (n=3), male 5xFAD (n=3) and male 5xFAD/*Neu1*^-/-^ (n=3) mice. **d**, Normalized expression of sialylated moPPCA in homogenized hippocampal tissue, relative to total protein levels displayed in *Extended Data Fig. 2d*. **e**, Quantification of β-hexosaminidase (β-Hex) in the CSF. **f**, Normalized expression of sialylated APP in homogenized hippocampal tissue, relative to total protein levels displayed in *Extended Data Fig. 2f*. **g**, Normalized expression of sialylated β-CTF in homogenized hippocampal tissue, relative to total protein levels displayed in *Extended Data Fig. 2g*. **h**, Quantification of Aβ42 concentrations in the CSF. **i**, Normalized expression of sialylated pro-TACE in homogenized hippocampal tissue, relative to total protein levels displayed in *Extended Data Fig. 2h*. **j**, Normalized expression of sialylated mature TACE in homogenized hippocampal tissue, relative to total protein levels displayed in *Extended Data Fig. 2i*. **k**, Normalized expression of sialylated BACE1 in homogenized hippocampal tissue, relative to total protein levels displayed in *Extended Data Fig. 2j*. **l**, Normalized expression of sialylated NCT in homogenized hippocampal tissue, relative to total protein levels displayed in *Extended Data Fig. 2k*. Actual p-values are given, any value less than or equal to 0.05 is considered significant.

Similar to AD patients, 5xFAD/*Neu1*^-/-^ mice exhibited higher levels of sialylated PPCA. However, unlike AD patients, these mice also showed elevated total PPCA levels, compared to 5xFAD controls, with these differences observed regardless of gender (Fig. 2d and Extended Data Fig. 2d). Within the 5xFAD background, female mice tended to have higher levels of total PPCA protein, regardless of Neu1 levels (Extended Data Fig. 2d). In addition, PPCA enzyme activity was generally higher in all Neu1-deficient animal, irrespective of their background, suggesting a link between increased sialylation of PPCA and enhanced enzyme activity (Extended Data Fig. 2e). We also confirmed that Neu1 deficiency led to an increase in extracellular β-Hex activity, indicating enhanced lysosomal exocytosis, as previously reported^35,38^. However, we now expanded on these findings by demonstrating that these significant increases occurred regardless of gender or background (Fig. 2e). Notably, under normal Neu1, β-Hex activity was higher in female 5xFAD mice than WT controls (Fig. 2e), suggesting an influence of the 5xFAD background on lysosomal biology in relation to Neu1. Additionally, female mice within the 5xFAD background consistently exhibited higher levels of β-Hex activity compared to males, implying a gender-specific difference in lysosomal function (Fig. 2e).

When characterizing APP, we found that both APP and β-CTF were elevated in Neu1-deficient mice. This effect was observed primarily in females within the 5xFAD background, regardless of Neu1 status (Extended Data Figs. 2f and 2g). However, Neu1 deficiency consistently increased the levels of sialylated APP and β-CTF, independent of total protein levels, genetic background, or sex. In addition, the 5xFAD background itself also consistently elevated these levels (Figs. 2f and 2g). Specifically, female 5xFAD mice always had higher levels of sialylated APP, irrespective of Neu1 levels, while male 5xFAD mice only exhibited higher sialylated β-CTF when Neu1 was deficient (Figs. 2f and 2g). We also confirmed that Neu1 deficiency led to increased release of Aβ42 (Fig. 2h), a process facilitated by excessive lysosomal exocytosis in these mice^38^. These findings suggest that the downregulation of NEU1 in AD patients may lead to increased release of toxic amyloid fragments, which are known to trigger neurotoxic microglia-mediated responses, thereby accelerating disease progression. Notably, when Neu1 was present, female 5xFAD mice released more Aβ42 than their male counterparts, but this effect was abolished when Neu1 was ablated (Fig. 2h). These results point to a gender bias that could help explain the greater plaque burden and faster disease progression observed in female mice.

Neu1 deficiency reduced the levels of both pro-and mature TACE, regardless of gender or background, which would effectively limit the non-amyloidogenic cleavage pathway in 5xFAD/*Neu1*^-/-^ mice (Extended Data Figs. 2h and 2i). Conversely, however, Neu1 deficiency increased the levels of sialylated pro-and mature TACE, regardless of gender or background (Figs. 2i and 2j). While Neu1 deficiency could elevate total BACE1 levels, this effect was less pronounced in the 5xFAD background, where no significant differences in BACE1 levels were observed between 5xFAD and 5xFAD/*Neu1*^-/-^ mice, regardless of gender (Extended Data Fig. 2j). However, the consistently higher BACE1 levels in the 5xFAD background combined with reduced TACE levels suggest a shift toward the pathogenic amyloidogenic cleavage pathway. Differential effects of Neu1 deficiency on sialylated BACE1 levels were observed across backgrounds. In *Neu1*^-/-^ mice, sialylated BACE1 levels were lower compared to wild-type (WT) mice. However, in the 5xFAD background, Neu1 deficiency led to an increase in sialylated BACE1 levels, with a more pronounced effect in females compared to males (Fig. 2k). With regard to NCT, Neu1 deficiency tended to decrease total NCT levels, regardless of gender or background (Extended Data Fig. 2k), but it had a unique effect in female 5xFAD/*Neu1*^-/-^ mice, where the levels of sialylated NCT increased (Fig. 2l). Taken altogether, these analyses of the sialylated protein levels of PPCA, APP, TACE, BACE1, and NCT suggest that these proteins are substrates of Neu1, and that their biochemical properties could be altered by the downregulation of NEU1, as seen in AD patients.

### NEU1 Knockout in 5xFAD Mice Alters the Turnover of APP, Its Secretases, and PPCA

Substrates of Neu1 exhibit altered turnover when Neu1 is deficient^35,39^. To confirm that our proteins of interests are substrates of Neu1, we assessed their turnover in hippocampal slice cultures treated with cycloheximide (CHX) for various periods of time. For PPCA, we observed slower turnover in *Neu1*^-/-^, 5xFAD, and 5xFAD/*Neu1*^-/-^ mice, regardless of gender (Fig. 3a). In females, the most notable difference in turnover rate occurred in 5xFAD/*Neu1*^-/-^ mice, where a clear delay in PPCA turnover was evident by 8 hours after CHX treatment (Fig. 3a). A similar delay was observed in male 5xFAD/*Neu1*^-/-^ mice. However, the most significant difference in turnover was in male 5xFAD mice, where PPCA initially accumulated until the 4-hour time point, after which the turnover accelerated and reached WT levels by the final 16-hour time point (Fig. 3a). The turnover patterns of PPCA were similar between males and females; however, Neu1 deficiency differentially impacted male and female mice. PPCA turnover was somewhat slower in male *Neu1*^-/-^ mice and much more significantly slower in male 5xFAD mice compared to their female counterparts (Extended Data Fig. 3a). Regarding APP, Neu1 deficiency tended to slow APP processing in the non-AD background but seemed to accelerate its turnover in the 5xFAD background, irrespective of gender (Fig. 3b). In 5xFAD mice, females processed APP slightly slower, while males showed significantly slower processing, even to the point of inhibition, given that APP accumulated over the CHX time course (Fig. 3b and Extended Data Fig. 3b). In 5xFAD/*Neu1*^-/-^ mice, males showed a slightly faster APP processing than controls, while females exhibited a more robust and excessive processing of APP (Fig. 3b and Extended Data Fig. 3b). The processing of β-CTF varied depending on background, gender, and Neu1 levels (Fig. 3c and Extended Data Fig. 3c). In females, β-CTF processing was slightly faster in *Neu1*^-/-^ mice, initially delayed but then accelerated in 5xFAD/*Neu1*^-/-^ mice, and remained initially unchanged before slowing down in 5xFAD mice, compared to WT controls (Fig. 3c). In males, β-CTF processing was slower in *Neu1*^-/-^, 5xFAD, and 5xFAD/*Neu1*^-/-^ mice compared to WT controls; however, Neu1 deficiency in the 5xFAD background enhanced β-CTF turnover (Fig. 3c).

**Fig. 3:**
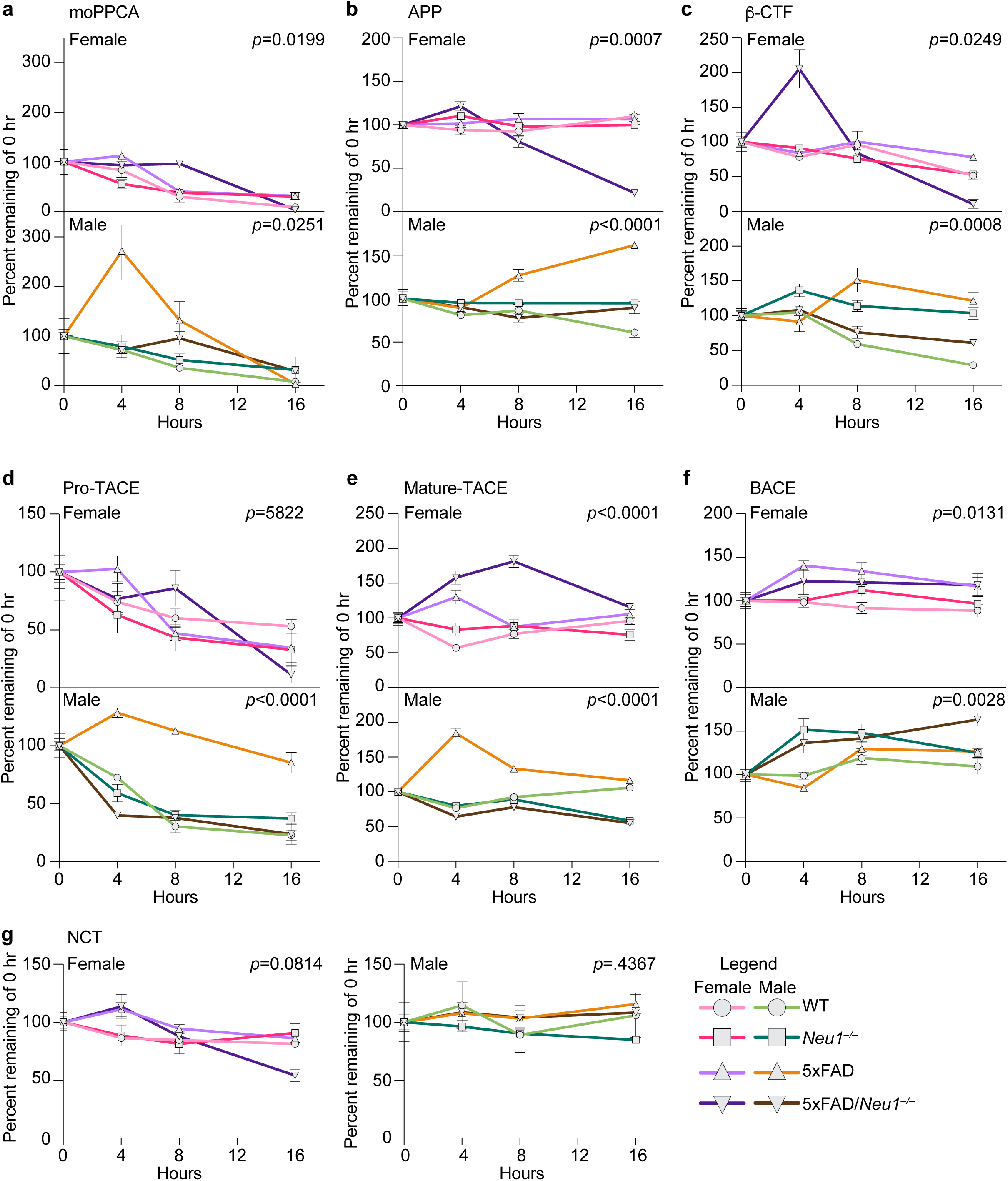
Turnover of proteins when Neu1 is deficient in 5xFAD mice. **a-g**, Measurements collected from hippocampal slice cultures of female WT (n=3), female *Neu1*^-/-^ (n=3), female 5xFAD (n=3), female 5xFAD/*Neu1*^-/-^ (n=3), male WT (n=3), male *Neu1*^-/-^ (n=3), male 5xFAD (n=3) and male 5xFAD/*Neu1*^-/-^ (n=3) mice treated with 100μM cycloheximide (CHX) for 0, 4, 8, and 16 h to calculate protein turnover, displayed as a percentage of quantified protein remaining compared to the 0 h timepoint, for **a**, moPPCA. **b**, APP. **c**, β-CTF. **d**, pro-TACE. **e**, mature TACE. **f**, BACE1. **g**, NCT. Female and male groups are plotted separately for comparisons between genetic backgrounds. Actual p-values are given, any value less than or equal to 0.05 is considered significant.

Neu1 deficiency also altered the turnover of the alpha and beta secretases. In female mice, Neu1 deficiency accelerated pro-TACE turnover in the non-AD background but slowed pro-TACE turnover in the 5xFAD background (Fig. 3d). In contrast, Neu1-deficient male mice showed accelerated pro-TACE turnover regardless of background (Fig. 3d). Notably, male 5xFAD mice exhibited significantly slower pro-TACE turnover, while 5xFAD/*Neu1*^-/-^ mice had enhanced turnover, compared to female counterparts. No gender differences were observed in *Neu1*^-/-^ mice (Extended Data Fig. 3d). For mature TACE, female mice in the 5xFAD background displayed slower TACE turnover, while male 5xFAD mice showed similar slow turnover; however, 5xFAD/*Neu1*^-/-^ males had slightly faster TACE turnover, compared to WT counterparts (Fig. 3e). When comparing genders, the turnover patterns of mature TACE followed a trend similar to pro-TACE, with male 5xFAD mice showing significantly slower turnover rates, while male 5xFAD/*Neu1*^-/-^ mice had accelerated turnover rates, compared to female counterparts (Extended Data Fig. 3e). Regarding BACE1, Neu1 deficiency slowed its turnover, regardless of gender or background (Fig. 3f). In the female population, 5xFAD mice exhibited the slowest BACE1 turnover, while, in males, turnover was slowest in 5xFAD/ *Neu1*^-/-^ and *Neu1*^-/-^ mice (Fig. 3f). When comparing genders, males showed slower BACE1 turnover, across background, except for 5xFAD mice, where female mice initially had slower turnover (Extended Data Fig. 3f). Lastly, Neu1 levels did not affect NCT turnover, though some trends were observed (Fig. 3g). NCT turnover was delayed in females and slowed/inhibited in males within the 5xFAD background (Fig. 3g). Gender comparison revealed that males generally processed NCT more slowly than female mice (Extended Data Fig. 3g). Overall, the CHX turnover analyses revealed that the proteolytic processing of PPCA, APP and the secretase proteins is influenced independently and differentially by the expression of the 5xFAD transgenes, the levels of Neu1 present, and the gender of the mouse. Combined with the sialylated protein level measurements, we can confirm that PPCA, APP, TACE, and BACE1 are substrates of Neu1. Although NCT appears to be influenced by loss of Neu1, whether it is a direct substrate remains unclear.

### Effects of PPCA overexpression in the 5xFAD model

It has been well established that NEU1 relies on PPCA for its full functionality^42,43^. Our findings now demonstrate that Neu1 levels heavily influence PPCA sialylation, turnover, and activity (Figs. 2 and 3). Furthermore, we have previously shown that injecting 5xFAD mice with viral vectors encoding human NEU1 and PPCA effectively reduces plaque burden, APP expression, and Aβ42 levels^38^. Therefore, the interaction between NEU1 and PPCA appears to play a crucial role in the extent and progression of AD pathology. To further investigate this, we aimed to determine which neural cell types were most dependent on the NEU1-PPCA interaction by generating and characterizing 5xFAD models with cell-specific overexpression of human PPCA (huPPCA).

### Under the CSF1R Promoter: Modulating Microglia and Monocyte Populations in 5xFAD Mice

We have reported earlier that transgenic mice overexpressing PPCA under the control of the CSFR1 promoter have restricted expression of the protein in the microglial/monocytic lineage^44^. When 5xFAD mice were crossed with these PPCA expressing transgenic mice (5xFAD/CSFR), no significant differences in plaque burden, plaque number, or average plaque size were detected compared to 5xFAD counterparts, regardless of gender (Figs. 4a-4a, and Extended Data Fig. 4a). The only noticeable difference in brain size was between male 5xFAD and 5xFAD/CSFR mice (Extended Data Fig. 4b). Interestingly, overexpression of PPCA had no significant impact on total brain Neu1 or PPCA enzyme activity (Extended Data Figs. 4c and 4d), although the total levels of mouse PPCA (moPPCA) and huPPCA were increased in 5xFAD/CSFR mice compared to controls, specifically 5xFAD and CSRF mice (Extended Data Figs. 4e and 4f). In contrast, the levels of sialylated moPPCA and huPPCA were reduced compared to controls (Fig. 4d and Extended Data Fig. 4g). These changes to PPCA were observed regardless of gender, although female 5xFAD/CSFR mice had higher levels of total moPPCA but lower levels of sialylated moPPCA compared to their male counterparts (Fig. 4d and Extended Data Fig. 4e). No differences were observed in the levels of huPPCA, whether total or sialylated, between male and female 5xFAD/CSFR mice (Extended Data Figs. 4f and 4g). These findings suggest that sialylated PPCA may be more enzymatically active than de-sialylated PPCA. Interestingly, we still measured a significantly higher extent of lysosomal exocytosis in 5xFAD/CSFR mice compared to 5xFAD mice, regardless of gender. Notably, female mice consistently displayed higher levels of extracellular β-Hex activity, irrespective of the background (Fig. 4e). This indicates that lysosomal exocytosis may still be contributing to the pathological development in these mice, possibly by releasing toxic Aβ42 peptides.

**Fig. 4:**
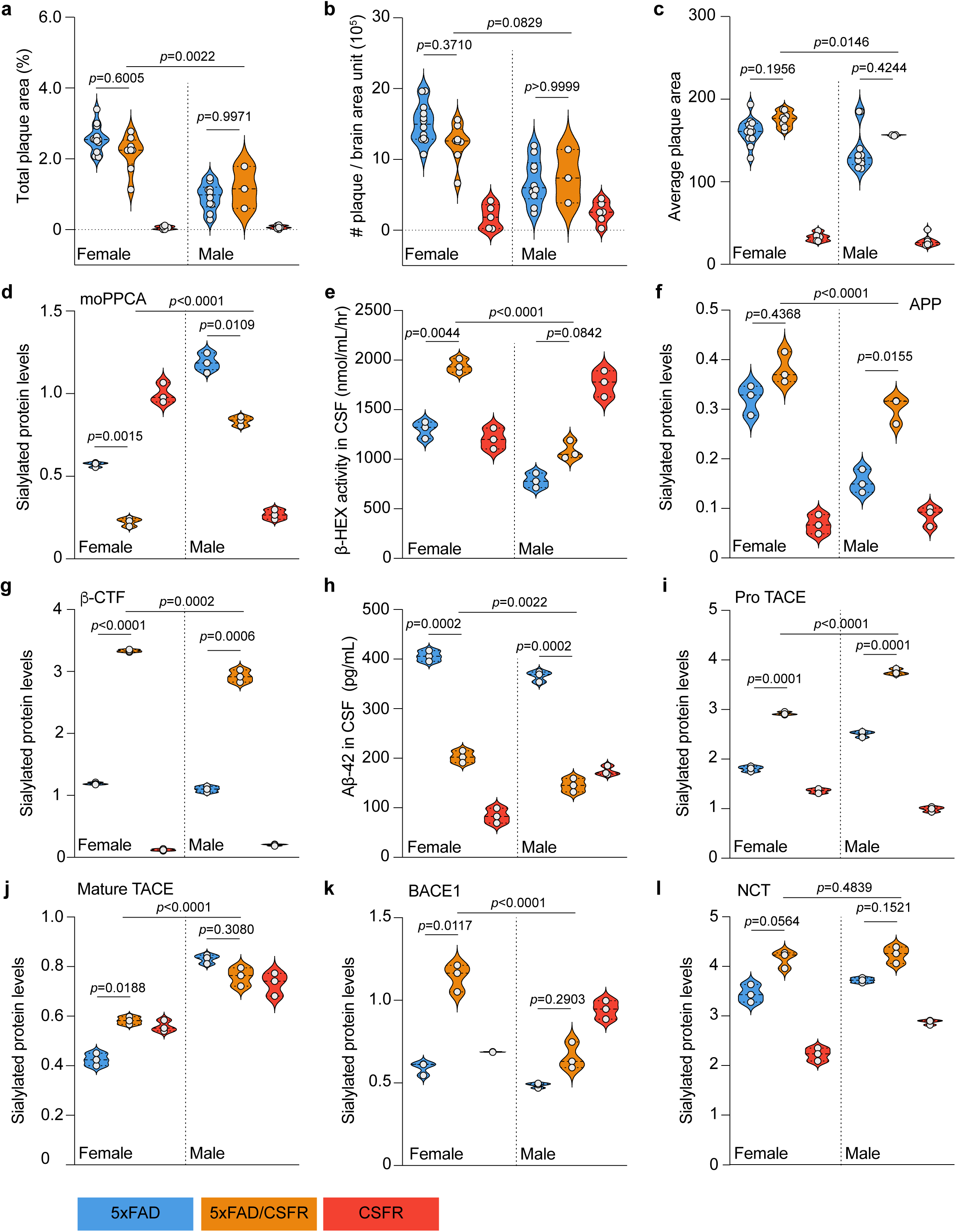
PPCA overexpression under the CSFR1 promoter in 5xFAD mice. **a-c**, Measurements collected from paraffin-embedded whole brain sections for female 5xFAD (n=12), female 5xFAD/CSFR (n=8), female CSFR (n=5), male 5xFAD (n=10), male 5xFAD/CSFR (n=3), and male CSFR (n=6) mice. **a**, Quantification of plaque burden, represented as amyloid^+^ (4G8^+^) area per section area, displayed as a percentage. **b**, Quantification of the average number of plaques per brain area unit (total number of plaques/total brain area). **c**, Quantification of the average plaque area (total plaque area/total plaque number). **d-l**, Measurements collected from female 5xFAD (n=3), female 5xFAD/CSFR (n=3), female CSFR (n=3), male 5xFAD (n=3), male 5xFAD/CSFR (n=3), and male CSFR (n=3) mice. **d**, Normalized expression of sialylated moPPCA in homogenized hippocampal tissue, relative to total protein levels displayed in *Extended Data Fig. 4e*. **e**, Quantification of β-hexosaminidase (β-Hex) in the CSF. **f**, Normalized expression of sialylated APP in homogenized hippocampal tissue, relative to total protein levels displayed in *Extended Data Fig. 4h*. **g**, Normalized expression of sialylated β-CTF in homogenized hippocampal tissue, relative to total protein levels displayed in *Extended Data Fig. 4i*. **h**, Quantification of Aβ42 concentrations in the CSF. **i**, Normalized expression of sialylated pro-TACE in homogenized hippocampal tissue, relative to total protein levels displayed in *Extended Data Fig. 4j*. **j**, Normalized expression of sialylated mature TACE in homogenized hippocampal tissue, relative to total protein levels displayed in *Extended Data Fig. 4k*. **k**, Normalized expression of sialylated BACE1 in homogenized hippocampal tissue, relative to total protein levels displayed in *Extended Data Fig. 4l*. **l**, Normalized expression of sialylated NCT in homogenized hippocampal tissue, relative to total protein levels displayed in *Extended Data Fig. 4m*. Actual p-values are given, any value less than or equal to 0.05 is considered significant.

Total levels of APP did not differ between 5xFAD and 5xFAD/CSFR mice, while β-CTF levels were reduced, regardless of gender (Extended Data Figs. 4h and 4i). This suggests that PPCA overexpression in the myeloid population may reduce amyloidogenic APP processing. However, sialylated levels of APP and β-CTF were increased in 5xFAD/CSFR mice compared to 5xFAD controls, irrespective of gender (Figs. 4f and 4f). In general, female 5xFAD/CSFR mice consistently exhibited higher levels of both total and sialylated APP and β-CTF compared to their male counterparts (Figs. 4f and 4g, and Extended Data Figs. 4h and 4i). Sialylation of APP has been shown to enhance its processing^23^, but when we measured the levels of Aβ42 in the CSF, we found that 5xFAD/CSFR mice released significantly less amyloid compared to 5xFAD controls, regardless of gender (Fig. 4h). This further suggests that amyloidogenic processing of APP is reduced in 5xFAD/CSFR mice. However, given that plaque burden remains unchanged between 5xFAD/CSFR and 5xFAD brains, it is possible that Aβ42 is retained within the brain parenchyma, rather than being filtered out into the CSF, which could explain the lower levels of Aβ42 detected in the CSF.

Amyloidogenic and non-amyloidogenic processing of APP is influence not only by the biochemical properties of APP but also by the biochemical properties of the secretases^21,24,26^. We found that total pro-TACE levels remained unchanged, while total mature TACE levels were higher in 5xFAD/CSFR mice compared to 5xFAD controls, regardless of gender (Extended Data Figs. 4j and 4k). Overall, total pro-and mature TACE levels were higher in female mice compared to male counterparts (Extended Data Figs. 4j and 4k). Additionally, levels of sialylated pro-TACE were higher in 5xFAD/CSFR mice compared to 5xFAD controls, irrespective of gender, with male mice consistently exhibiting higher levels than their female counterparts (Fig. 4i). This pattern slightly differed for mature TACE, where levels of sialylated mature TACE were still higher in female 5xFAD/CSFR mice but were lower in male 5xFAD/CSFR mice compared to 5xFAD controls. However, levels remained always higher in male mice compared to their female counterparts (Fig. 4j). Total levels of BACE1 were unchanged between female 5xFAD/CSFR and 5xFAD mice but were lower in male 5xFAD/CSFR mice compared to 5xFAD controls (Extended Data Fig. 4l). Contrary to this, levels of sialylated BACE1 were higher in 5xFAD/CSFR mice compared to 5xFAD controls, regardless of gender, with levels in females being higher compared to males (Fig. 4k). Total levels of NCT mirrored BACE1, showing no change between female 5xFAD/CSFR and 5xFAD mice but lower levels in male 5xFAD/CSFR mice (Extended Data Fig. 4m). Sialylated NCT levels were consistently higher in 5xFAD/CSFR mice compared to 5xFAD controls, regardless of gender (Fig. 4l). Altogether, these data demonstrate that overexpression of huPPCA protein in microglia influences the biochemical properties of secretase enzymes and subunits but has little impact on the plaque pathology that develops in the brain parenchyma.

### Under the NSE Promoter: Modulating the Neuronal Population in 5xFAD Mice

When we crossed 5xFAD mice with mice overexpressing PPCA under the NSE promoter (5xFAD/NSE), we observed a greater effect in females than in males for all plaque characterization measurements, compared to 5xFAD controls (Figs. 5a-5c and Extended Data Fig. 5a). There was no difference in total brain areas between genders (Extended Data Fig. 5b). In terms of total plaque area, female 5xFAD/NSE mice showed a marked reduction in DAB staining, while male mice showed only a slight decrease compared to 5xFAD controls (Fig. 5a). For the average number of plaques per brain area unit, we found that female 5xFAD/NSE mice tended to have fewer plaques compared to 5xFAD controls, while no difference was observed in males (Fig. 5b). Regarding average plaque size, we measured a slight reduction in 5xFAD/NSE mice compared to 5xFAD controls, regardless of gender (Fig. 5c). Overall, overexpressing PPCA under the NSE promoter effectively reduced plaque pathology in 5xFAD mice. Interestingly, while PPCA overexpression did not obviously alter PPCA enzyme activity levels, it appeared to increase Neu1 activity (Extended Data Figs. 5c and 5d). Furthermore, the total PPCA protein levels were significantly higher; moPPCA expression was higher in 5xFAD/NSE mice compared to 5xFAD controls and huPPCA expression was similarly higher in 5xFAD/NSE mice compared to NSE controls, regardless of gender (Extended Data Figs. 5e and 5f). Female 5xFAD/NSE mice consistently showed the highest total PPCA protein levels. Similar to the observations in 5xFAD/CSFR mice, sialylated moPPCA and huPPCA levels were lower in 5xFAD/NSE mice compared to controls, regardless of gender; however, these differences were less pronounced than those observed in 5xFAD/CSFR mice (Fig. 5d and Extended Data Fig. 5g). Interestingly, β-Hex enzyme activity in the CSF was only slightly increased in female 5xFAD/NSE mice, while it was significantly higher in male 5xFAD/NSE mice, compared to 5xFAD controls (Fig. 5e). These variations could account for the more pronounced effect on plaque pathology seen in female 5xFAD/NSE mice, as compared to males, given the correlation between lysosomal exocytosis and amyloid release.

**Fig. 5:**
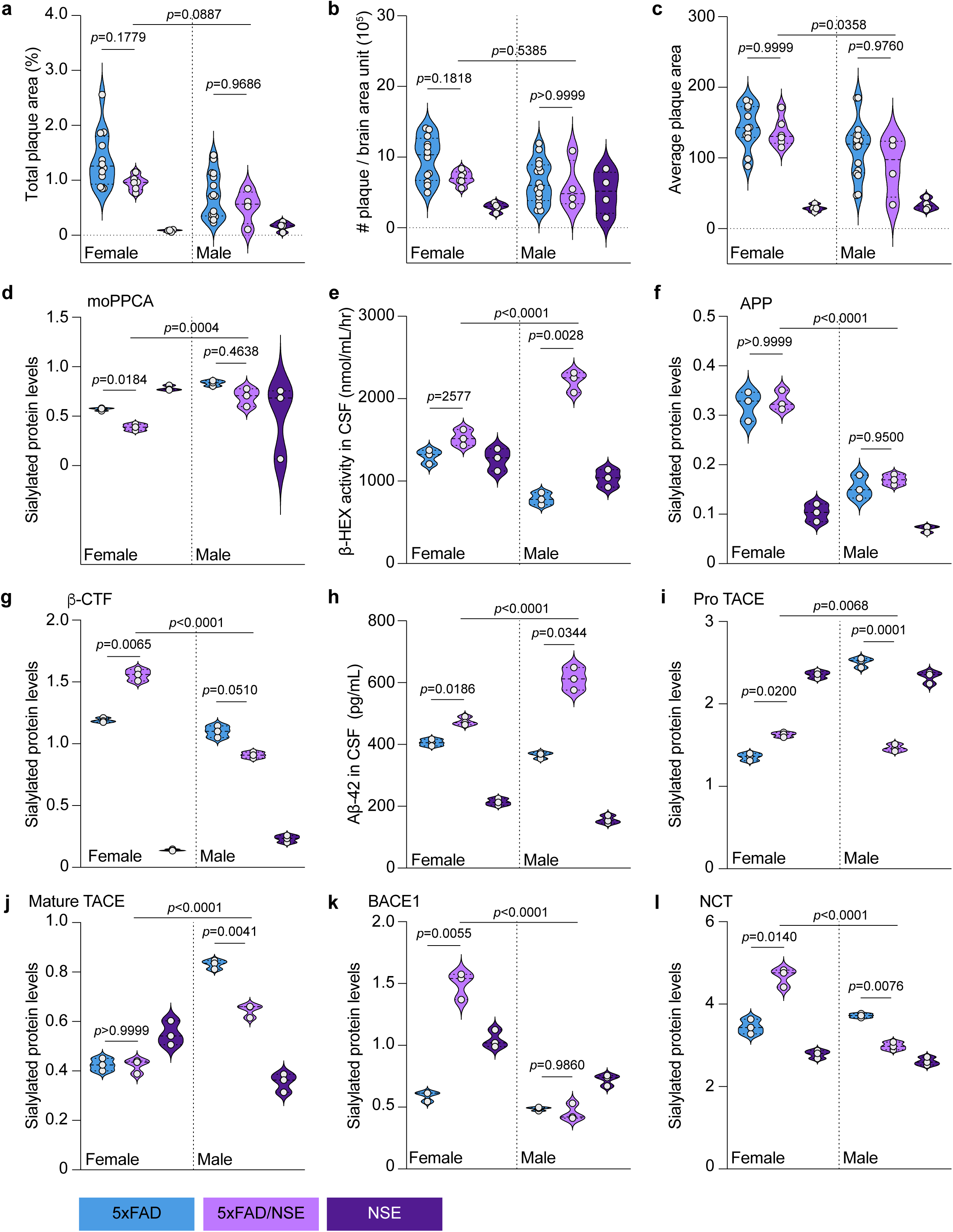
PPCA overexpression under the NSE promoter in 5xFAD mice. **a-c**, Measurements collected from paraffin-embedded whole brain sections for female 5xFAD (n=12), female 5xFAD/NSE (n=6), female NSE (n=4), male 5xFAD (n=16), male 5xFAD/NSE (n=4), and male NSE (n=4) mice. **a**, Quantification of plaque burden, represented as amyloid^+^ (4G8^+^) area per section area, displayed as a percentage. **b**, Quantification of the average number of plaques per brain area unit (total number of plaques/total brain area). **c**, Quantification of the average plaque area (total plaque area/total plaque number). **d-l**, Measurements collected from female 5xFAD (n=3), female 5xFAD/NSE (n=3), female NSE (n=3), male 5xFAD (n=3), male 5xFAD/NSE (n=3), and male NSE (n=3) mice. **d**, Normalized expression of sialylated moPPCA in homogenized hippocampal tissue, relative to total protein levels displayed in *Extended Data Fig. 5e*. **e**, Quantification of β-hexosaminidase (β-Hex) in the CSF. **f**, Normalized expression of sialylated APP in homogenized hippocampal tissue, relative to total protein levels displayed in *Extended Data Fig. 5h*. **g**, Normalized expression of sialylated β-CTF in homogenized hippocampal tissue, relative to total protein levels displayed in *Extended Data Fig. 5i*. **h**, Quantification of Aβ42 concentrations in the CSF. **i**, Normalized expression of sialylated pro-TACE in homogenized hippocampal tissue, relative to total protein levels displayed in *Extended Data Fig. 5j*. **j**, Normalized expression of sialylated mature TACE in homogenized hippocampal tissue, relative to total protein levels displayed in *Extended Data Fig. 5k*. **k**, Normalized expression of sialylated BACE1 in homogenized hippocampal tissue, relative to total protein levels displayed in *Extended Data Fig. 5l*. **l**, Normalized expression of sialylated NCT in homogenized hippocampal tissue, relative to total protein levels displayed in *Extended Data Fig. 5m*. Actual p-values are given, any value less than or equal to 0.05 is considered significant.

Total APP protein expression was enhanced in female 5xFAD/NSE mice, with no differences observed in males when compared to 5xFAD controls (Extended Data Fig. 5h). A similar trend was seen for β-CTF, with higher total levels in female 5xFAD/NSE mice, but lower levels in males, compared to 5xFAD controls (Extended Data Fig. 5i). Interestingly, sialylated APP levels did not differ between 5xFAD/NSE and 5xFAD mice, regardless of gender. However, sialylated β-CTF levels were higher in female 5XFAD/NSE mice and lower in male mice, compared to controls (Figs. 5f and 5g). This pattern contrasted with 5xFAD/CSFR mice, which displayed drastically higher levels of sialylated β-CTF compared to controls, potentially explaining the reduced plaque pathology seen in female 5xFAD/NSE mice. Likewise, the amount of Aβ42 released into the CSF differed significantly across mouse models. While Aβ42 release was significantly reduced in 5xFAD/CSFR mice, Aβ42 concentrations were significantly higher in the CSF of 5xFAD/NSE mice compared to 5xFAD (Fig. 5h). These findings suggest a relationship between the level of sialylation on β-CTF and the amount of Aβ42 released into the CSF.

We observed distinct gender differences in secretase levels between 5xFAD/NSE mice, which were not as apparent in 5xFAD/CSFR mice. These differences may also contribute to the variations in plaque pathology. Total levels of pro-TACE did not significantly differ between the sexes in 5xFAD/NSE mice, but mature TACE levels were higher, regardless of gender, compared to controls (Extended Data Figs. 5j and 5k). In comparison to 5xFAD mice, female 5xFAD/NSE mice showed increased levels of sialylated pro-TACE, while males showed a decrease. Additionally, the levels of sialylated mature TACE remained unchanged in female 5xFAD/NSE mice but were decreased in males (Figs. 5i and 5j). Total levels of BACE1 were lower in both male and female 5xFAD/NSE mice, although sialylated BACE1 levels were increased in female 5xFAD/NSE mice and unchanged in males, compared to controls (Fig. 5k and Extended Data Fig. 5l). For NCT, total levels did not differ between female 5xFAD and 5xFAD/NSE mice but were lower in male 5xFAD/NSE mice. Sialylated NCT levels were higher in female 5xFAD/NSE mice and lower in males, compared to 5xFAD controls (Fig. 5l and Extended Data Fig. 5m). Reflecting on our findings from Fig. 1, where increased sialylation of the secretases or their components correlated with higher enzyme activity, we can speculate that one reason for the altered plaque pathology observed in female 5xFAD/NSE mice, but not in males, could be due to changes in the enzyme activity of the secretases. Specifically, the reduced sialylation of mature TACE, but not BACE1, may lead to decreased non-amyloidogenic activity, while maintaining amyloidogenic activity, thereby promoting the processing of APP to produce similar plaque pathology between male 5xFAD/NSE and 5xFAD mice.

### Under the GFAP Promoter: Modulating Astrocyte and Neuronal Populations in 5xFAD Mice

We previously reported that transgenic mice overexpressing PPCA under the control of the GFAP promoter exhibit expression of the protein in astrocytes, with occasional ectopic expression in some neurons^45^. When these transgenic mice were crossed into the 5xFAD line to generate 5xFAD/GFAP mice, we found no significant differences between 5xFAD and 5xFAD/GFAP animals in total plaque burden, average plaque number per brain area, average plaque size, or total brain area, irrespective of sex (Figs. 6a-6c and Extended Data Figs. 6a and 6b). However, female mice tended to display higher plaque burdens, a greater number of plaques per area, and larger plaque sizes compared to their male counterparts within the same genotype (Figs. 6a-6c). Interestingly, the enzymatic activity levels of both Neu1 and PPCA were elevated in 5xFAD/GFAP mice relative to 5xFAD controls (Extended Data Figs. 6c and 6d). This increase was accompanied by higher levels of moPPCA in 5xFAD/GFAP mice, while levels of huPPCA were reduced when compared to GFAP transgenic controls (Extended Data Figs. 6e and 6f). Notably, female mice consistently exhibited higher expression levels of PPCA than their male counterparts. The increased Neu1 activity likely contributed to the significantly reduced levels of sialylated moPPCA and huPPCA observed in 5xFAD/GFAP mice relative to the appropriate controls (Fig. 6d and Extended Data Fig. 6g). However, the differences in Neu1 activity alone cannot fully explain the pronounced gender-specific variations observed in β-Hex enzyme activity, used to monitor the extent of lysosomal exocytosis. Extracellular β-Hex activity in the CSF was decreased in female 5xFAD/GFAP mice but increased in male 5xFAD/GFAP mice, compared to their respective 5xFAD controls (Fig. 6e). Within the same genotype group, female 5xFAD mice hade higher β-Hex activity than males. However, overexpression of PPCA under the GFAP promoter resulted in elevated β-Hex activity in male mice, regardless of the presence or absence of the 5xFAD transgene (Fig. 6e).

**Fig. 6:**
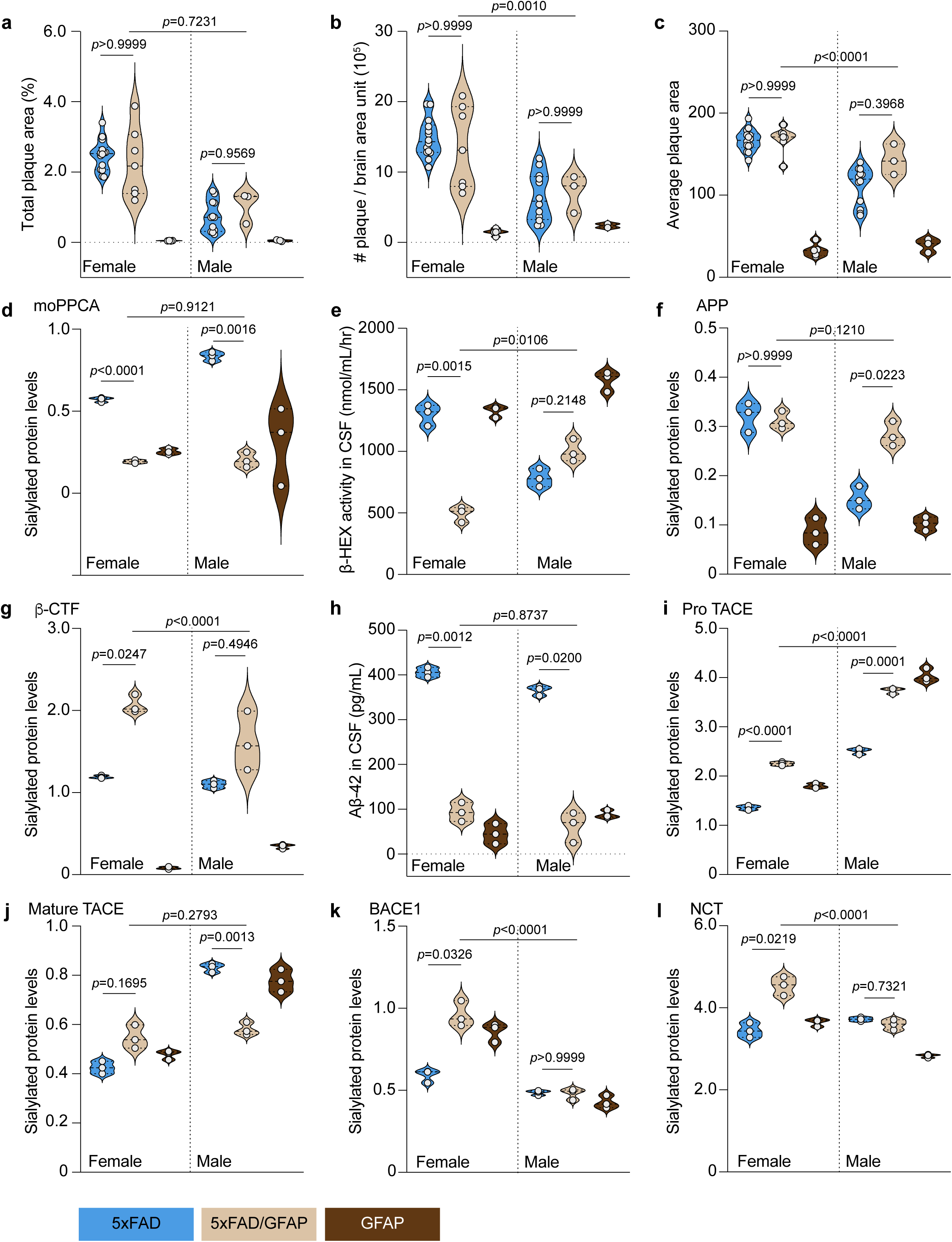
PPCA overexpression under the GFAP promoter in 5xFAD mice. **a-c**, Measurements collected from paraffin-embedded whole brain sections for female 5xFAD (n=14), female 5xFAD/GFAP (n=7), female GFAP (n=5), male 5xFAD (n=12), male 5xFAD/GFAP (n=3), and male GFAP (n=3) mice. **a**, Quantification of plaque burden, represented as amyloid^+^ (4G8^+^) area per section area, displayed as a percentage. **b**, Quantification of the average number of plaques per brain area unit (total number of plaques/total brain area). **c**, Quantification of the average plaque area (total plaque area/total plaque number). **d-l**, Measurements collected from female 5xFAD (n=3), female 5xFAD/GFAP (n=3), female GFAP (n=3), male 5xFAD (n=3), male 5xFAD/GFAP (n=3), and male GFAP (n=3) mice. **d**, Normalized expression of sialylated moPPCA in homogenized hippocampal tissue, relative to total protein levels displayed in *Extended Data Fig. 6e*. **e**, Quantification of β-hexosaminidase (β-Hex) in the CSF. **f**, Normalized expression of sialylated APP in homogenized hippocampal tissue, relative to total protein levels displayed in *Extended Data Fig. 6h*. **g**, Normalized expression of sialylated β-CTF in homogenized hippocampal tissue, relative to total protein levels displayed in *Extended Data Fig. 6i*. **h**, Quantification of Aβ42 concentrations in the CSF. **i**, Normalized expression of sialylated pro-TACE in homogenized hippocampal tissue, relative to total protein levels displayed in *Extended Data Fig. 6j*. **j**, Normalized expression of sialylated mature TACE in homogenized hippocampal tissue, relative to total protein levels displayed in *Extended Data Fig. 6k*. **k**, Normalized expression of sialylated BACE1 in homogenized hippocampal tissue, relative to total protein levels displayed in *Extended Data Fig. 6l*. **l**, Normalized expression of sialylated NCT in homogenized hippocampal tissue, relative to total protein levels displayed in *Extended Data Fig*. 6m. Actual p-values are given, any value less than or equal to 0.05 is considered significant.

Total APP protein levels followed a similar pattern between male and female mice, although expression was consistently higher in females; 5xFAD/GFAP mice had slightly reduced total APP compared to 5xFAD controls (Extended Data Fig. 6h). However, sialylated APP levels showed a gender-dependent difference. While females consistently had higher sialylated APP levels overall, there was no significant difference between female 5xFAD/GFAP and 5xFAD mice. In contrast, sialylated APP levels were significantly higher in male 5xFAD/GFAP mice compared to male 5XFAD controls (Fig. 6f). Regarding β-CTF, total levels were reduced in 5xFAD/GFAP mice compared to 5xFAD mice, across both sexes. However, sialylated β-CTF levels were significantly higher in female 5xFAD/GFAP mice and tended to be elevated in males as well, relative to controls (Fig. 6g and Extended Data Fig. 6i). Similar to what was observed in 5xFAD/CSFR mice, Aβ42 concentrations in the CSF were significantly lower in 5xFAD/GFAP mice compared to 5xFAD controls, regardless of sex (Fig. 6h). Remarkably, 5xFAD/CSFR and 5xFAD/GFAP mice displayed similar plaque morphology and comparable biochemical profiles of APP family proteins, setting them apart from 5sFAD/NSE mice. These findings suggest potential shared roles for microglia and astrocytes in modulating APP processing, in contrast to the distinct effects observed in neurons.

However, this pattern did not extend to the characterization of secretase proteins in the 5xFAD/GFAP model. Regarding TACE, total levels of the pro-form were similar between 5xFAD/GFAP and 5xFAD mice, while levels of the mature form were significantly higher in 5xFAD/GFAP mice, irrespective of sex. Females tended to have higher levels of both pro-and mature TACE compared to males (Extended Data Figs. 6j and 6k). In contrast, levels of sialylated pro-TACE were elevated in 5xFAD/GFAP mice relative to 5xFAD controls, with males showing higher levels than females (Fig. 6i). For the mature form, sialylated TACE levels tended to be higher in female 5xFAD/GFAP mice but were significantly reduced in male mice, compared to their respective 5xFAD controls (Fig. 6j). Notably, this gender difference resulted in comparable levels of sialylated mature TACE between female and male 5xFAD/GFAP mice, while male 5xFAD mice exhibited significantly higher levels than females (Fig. 6j). As for BACE1, total protein levels were decreased in female 5xFAD/GFAP mice compared to 5xFAD controls, while no significant differences were observed in males (Extended Data Fig. 6l). Among 5XFAD mice, no gender differences were observed in total BACE1 levels. However, in the 5xFAD/GFAP group, females had lower total BACE1 levels compared to males (Extended Data Fig. 6l). Notably, the pattern for sialylated BACE1 were nearly opposite: levels were elevated in female 5xFAD/GFAP mice but remained unchanged in males, when compared to their respective 5xFAD controls (Fig. 6k). Additionally, while there was no difference in sialylated BACE1 levels between female and male 5xFAD mice, female 5xFAD/GFAP displayed significantly higher levels than their male counterparts (Fig. 6k). Total levels of NCT were comparable between 5xFAD/GFAP and 5xFAD mice, with no significant differences between sexes within the same genotype (Extended Data Fig. 6m). However, sialylated NCT levels were highest in female 5xFAD/GFAP mice across all comparisons (Fig. 6l).

These findings indicate that the biochemical characteristics of secretase proteins in 5xFAD/GFAP mice more closely resemble those seen in the 5xFAD/NSE mice, while differing markedly from the 5xFAD/CSFR mice. This may be attributed to the presence of neuronal PPCA expression in both NSE and GFAP transgenic models, underscoring the distinct roles that specific cell types, particularly neurons, play distinct roles in APP processing.

### Overexpression of NEU1 and PPCA Under the GFAP Promoter

Thus far, PPCA overexpression has been shown to ameliorate certain aspects of AD pathology. However, considering the functional interdependence between Neu1 and PPCA, these beneficial effects may be offset by disproportionate increases in Neu1 expressions. To explore this possibility, we generated mice overexpressing both PPCA and NEU1 under the GFAP promoter (5xFAD/GNIP) and analyzed their brains pathology. These mice exhibited some similarities with other PPCA-overexpression models but also displayed notable distinctions from Neu1-deficient 5xFAD mice, resulting in a unique biochemical and pathological profile. In female 5xFAD/GNIP mice, total plaque burden was significantly reduced compared to 5xFAD controls, while no differences were observed between males of the two genotypes (Fig. 7a and Extended Data Fig. 7a), a contrast to findings in 5xFAD/*Neu1*^-/-^ mice. Regarding plaque density, measured as the average number of plaques per unit brain area, no significant differences were found between 5xFAD/GNIP and 5xFAD mice overall. However, within each genotype, females consistently exhibited a higher plaque count than their male counterparts (Fig. 7b). Consistent with the reduction in total plaque burden, average plaque size was significantly reduced in female 5xFAD/GNIP mice compared to 5xFAD controls, while no difference was observed in males (Fig. 7c). Total brain size remained unchanged across all groups (Extended Data Fig. 7b). Similar to what was seen in 5xFAD/NSE mice, 5xFAD/GNIP mice had increased levels of Neu1 enzyme activity but showed no change in PPCA activity (Extended Data Figs. 7c and 7d). The absence of increased PPCA activity may be explained by the opposing trends in PPCA protein levels: 5xFAD/GNIP mice had higher levels of moPPCA but lower levels of huPPCA, compared to 5xFAD controls, regardless of gender (Extended Data Figs. 7e and 7f). in line with observations in

**Fig. 7:**
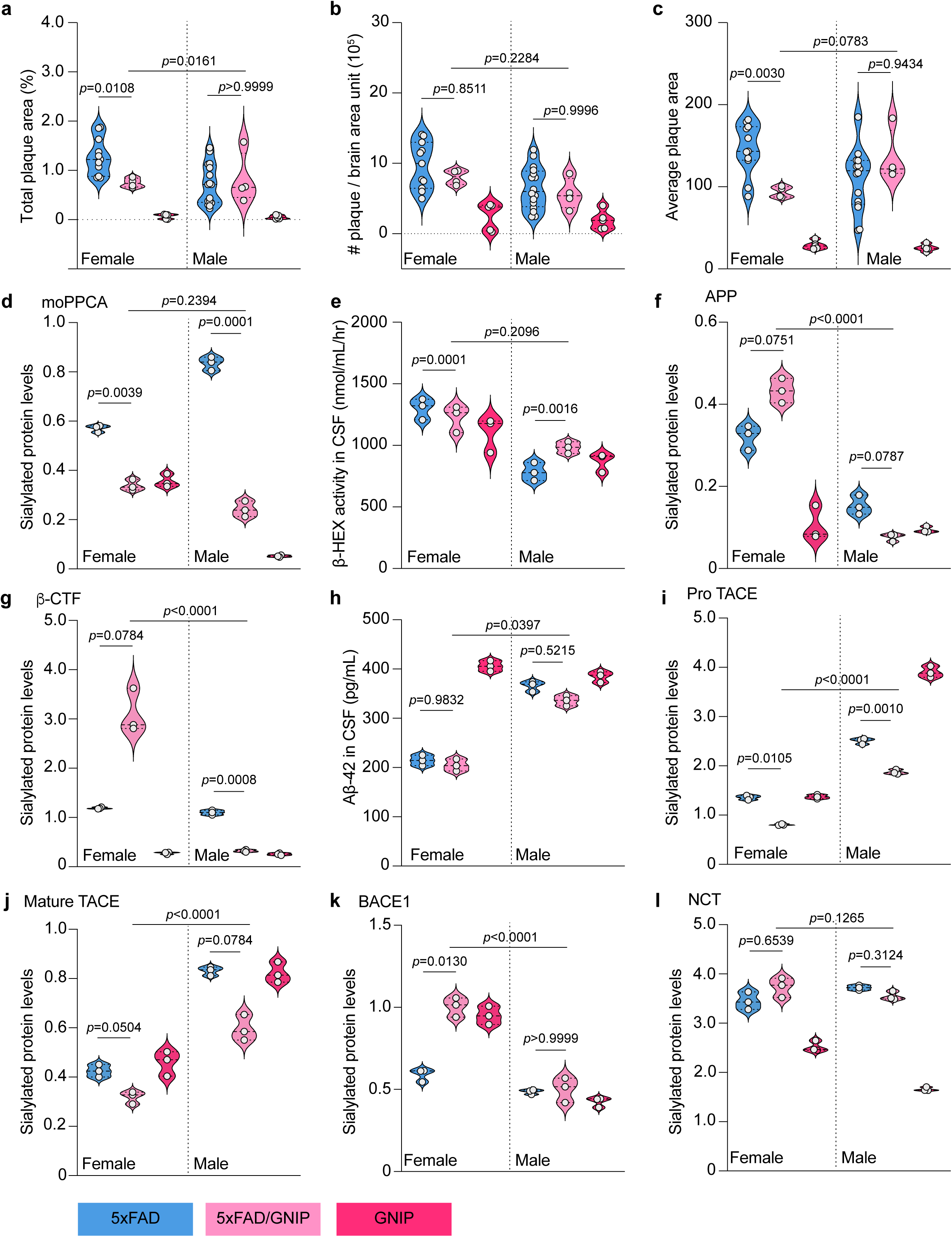
PPCA and NEU1 overexpression under the GFAP promoter in 5xFAD mice. **a-c**, Measurements collected from paraffin-embedded whole brain sections for female 5xFAD (n=11), female 5xFAD/GNIP (n=4), female GNIP (n=5), male 5xFAD (n=16), male 5xFAD/GNIP (n=4), and male GNIP (n=5) mice. **a**, Quantification of plaque burden, represented as amyloid^+^ (4G8^+^) area per section area, displayed as a percentage. **b**, Quantification of the average number of plaques per brain area unit (total number of plaques/total brain area). **c**, Quantification of the average plaque area (total plaque area/total plaque number). **d-l**, Measurements collected from female 5xFAD (n=3), female 5xFAD/GNIP (n=3), female GNIP (n=3), male 5xFAD (n=3), male 5xFAD/GNIP (n=3), and male GNIP (n=3) mice. **d**, Normalized expression of sialylated moPPCA in homogenized hippocampal tissue, relative to total protein levels displayed in *Extended Data Fig. 7e*. **e**, Quantification of β-hexosaminidase (β-Hex) in the CSF. **f**, Normalized expression of sialylated APP in homogenized hippocampal tissue, relative to total protein levels displayed in *Extended Data Fig. 7h*. **g**, Normalized expression of sialylated β-CTF in homogenized hippocampal tissue, relative to total protein levels displayed in *Extended Data Fig. 7i*. **h**, Quantification of Aβ42 concentrations in the CSF. **i**, Normalized expression of sialylated pro-TACE in homogenized hippocampal tissue, relative to total protein levels displayed in *Extended Data Fig. 7j*. **j**, Normalized expression of sialylated mature TACE in homogenized hippocampal tissue, relative to total protein levels displayed in *Extended Data Fig. 7k*. **k**, Normalized expression of sialylated BACE1 in homogenized hippocampal tissue, relative to total protein levels displayed in *Extended Data Fig. 7l*. **l**, Normalized expression of sialylated NCT in homogenized hippocampal tissue, relative to total protein levels displayed in *Extended Data Fig. 7m*. Actual p-values are given, any value less than or equal to 0.05 is considered significant.

5xFAD/CSFR and 5xFAD/GFAP mice, both sialylated moPPCA and huPPCA levels were significantly reduced in 5xFAD/GNIP mice relative to controls (Fig. 7d and Extended Data Fig. 7g). However, a unique facet of 5xFAD/GNIP mice was the lack of significant changes in extracellular β-Hex activity compared to 5xFAD controls (Fig. 7e). Taken together, these results suggest that co-overexpression of NEU1 and PPCA yields more favorable outcomes than overexpression of PPCA alone, highlighting the importance of maintaining a functional balance between these two interdependent enzymes.

APP was also uniquely affected in the 5xFAD/GNIP model. Total APP levels were elevated in female 5xFAD/GNIP mice but remained unchanged in males compared to 5xFAD controls (Extended Data Fig. 7h). Sialylated APP levels followed a similar trend – tending to be higher in females 5xFAD/GNIP mice but significantly reduced in males relative to controls (Fig. 7f). Downstream of APP, total β-CTF levels did not differ between 5xFAD/GNIP and 5xFAD mice in either sex. However, sialylated β-CTF levels were modestly increased in female 5xFAD/GNIP mice and significantly decreased in males compared to their respective controls (Fig. 7g and Extended Data Fig. 7i). Remarkably, 5xFAD/GNIP mice were the only transgenic model in this study that did not show any differences in CSF Aβ42 concentrations relative to 5xFAD mice, regardless of gender (Fig. 7h). These unique and distinct changes in the expression and sialylation patterns of APP family of proteins may underlie the observed significant reduction in plaque pathology in female 5xFAD/GNIP mice.

Analysis of the secretase proteins in 5xFAD/GNIP mice, in comparison to the other transgenic models, revealed distinct patterns that correlated with Neu1 activity levels and total moPPCA levels. Total levels of pro-TACE and mature TACE were largely unchanged between 5xFAD/GNIP and 5xFAD mice, with the exception of a significant reduction in mature TACE levels in female 5xFAD/GNIP mice (Extended Data Figs. 7j and 7k). In contrast to 5xFAD/*Neu1*^-/-^, 5xFAD/CSFR, and 5xFAD/GFAP mice, but closely resembling the 5xFAD/NSE model, levels of sialylated pro-TACE and mature TACE were significantly reduced in 5xFAD/GNIP mice compared to 5xFAD controls, regardless of gender (Figs. 7i and 7j). Both 5xFAD/NSE and 5xFAD/GNIP mice shared the distinctive feature of elevated Neu1 activity without a corresponding increase in PPCA activity, suggesting that increased Neu1 activity alone is sufficient to drive a reduction in the sialylation of TACE proteins. In terms of BACE1, total levels were decreased in female 5xFAD/GNIP mice, but increased in males, relative to 5XFAD controls (Extended Data Fig. 7l). Consistent with all other models characterized, levels of sialylated BACE1 were increased in female 5xFAD/GNIP mice but remained unchanged in males compared to 5xFAD controls (Fig. 7k). Another common feature across all models was an increase in the total moPPCA levels. Together these findings suggest that PPCA expression may influence the levels of sialylated BACE1, potentially in a sex-dependent manner. Total levels of NCT were slightly reduced in 5xFAD/GNIP mice, while sialylated NCT levels were unchanged compared to 5xFAD controls, regardless of gender (Fig. 7l and Extended Data Fig. 7m). These results present a more complex picture. Although both Neu1 enzyme activity and moPPCA protein levels appear to influence NCT, the relationship is not straightforward and remains difficult to interpret within the context of these current models.

### Impact of NEU1 and PPCA Levels on Secretase Enzymatic Activity

Until now, we chose to characterize each model individually with regards to plaque pathology, Neu1 and PPCA activity, and the biochemical properties of PPCA, APP, β-CTF, pro TACE, mature TACE, BACE1, and NCT. We now decided to present the combined analysis of alpha-, beta-and gamma-secretase activity to facilitate a comparison between the models and provide a final assessment of the impact of Neu1 and PPCA on the development of AD pathology in 5xFAD mice. We found that secretase activity was consistently higher in female 5xFAD mice compared to their male counterparts (Extended Data Figs. 8a-8c). This difference could partly explain the mark gender disparity in AD pathology, with female mice exhibiting a more severe phenotype. However, this trend did not hold true for our genetic models.

**Fig. 8:**
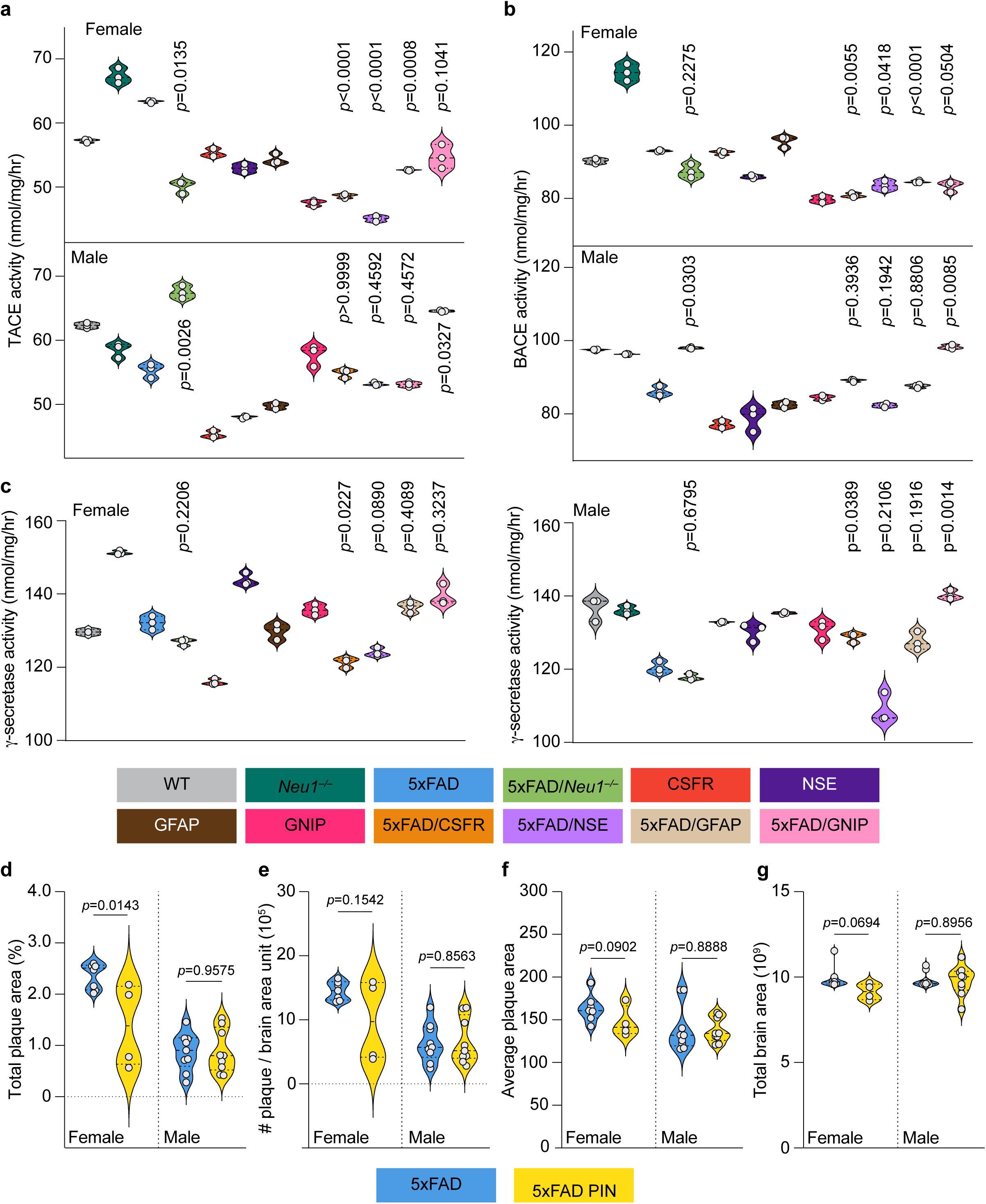
Secretase enzyme activities across the 5xFAD models and AAV-PIN injections in 5xFAD mice. **a-c**, Measurements collected from homogenized hippocampal tissue from female and male WT, *Neu1*^-/-^, 5xFAD, 5xFAD/*Neu1*^-/-^, CSFR, NSE, GFAP, GNIP, 5xFAD/CSFR, 5xFAD/NSE, 5xFAD/GFAP, and 5xFAD/GNIP mice (n=3 for all). Female and male groups are plotted separately for comparisons between genetic backgrounds. Displayed p-values are calculated against 5xFAD samples. **a**, Quantification of TACE enzyme activity. **b**, Quantification of BACE1 enzyme activity. **c**, Quantification of γ-secretase enzyme activity. **d-g**, Measurements collected from paraffin-embedded whole brain sections for female 5xFAD (n=7), female 5xFAD/AAV-PIN (n=4), male 5xFAD (n=9) and male 5xFAD/AAV-PIN (n=9). **d**, Quantification of plaque burden, represented as amyloid^+^ (4G8^+^) area per section area, displayed as a percentage. **e**, Quantification of the average number of plaques per brain area unit (total number of plaques/total brain area). **f**, Quantification of the average plaque area (total plaque area/total plaque number). **g**, Quantification of total brain area. Actual p-values are given, any value less than or equal to 0.05 is considered significant.

For alpha-secretase, female 5xFAD mice consistently had higher enzyme activity compared to our female genetic models, 5xFAD/*Neu1*^-/-^, 5xFAD/CSFR, 5xFAD/NSE, 5xFAD/GFAP and 5xFAD/GNIP (Fig. 8a). In contrast, the male mice showed more variability; alpha-secretase activity was increased in 5xFAD/*Neu1*^-/-^ and 5xFAD/GNIP mice, slightly decreased in male 5xFAD/NSE and 5xFAD/GFAP mice, and unchanged in 5xFAD/CSFR mice compared to 5xFAD controls (Fig. 8a). When comparing the male and female populations within our genetic models, the male populations generally had higher alpha-secretase enzyme activity, except for 5xFAD/GFAP, where no gender difference was observed (Extended Data Fig. 8a). Among all our 5xFAD models, male 5xFAD/*Neu1*^-/-^ mice, followed closely by male 5xFAD/GNIP mice, had the highest alpha-secretase activity, while female 5xFAD/NSE exhibited the lowest activity levels (Extended Data Fig. 8a).

Beta-secretase followed a pattern similar to that of alpha-secretase. In the female population, enzyme activity was reduced across all of our genetic models, 5xFAD/*Neu1*^-/-^, 5xFAD/CSFR, 5xFAD/NSE, 5xFAD/GFAP and 5xFAD/GNIP, compared to 5xFAD controls (Fig. 8b). In contrast, beta-secretase activity increased in male 5xFAD/*Neu1*^-/-^ and 5xFAD/GNIP mice, remained unchanged in male 5xFAD/CSFR and 5xFAD/GFAP mice and increased in 5xFAD/NSE mice compared to 5xFAD controls (Fig. 8b). When comparing males to females, the male population consistently had higher enzyme activity across all genetic models, with the exception of the 5xFAD/NSE group, where no gender differences were recorded (Extended Data Fig. 8b). Among all the 5xFAD models, male 5xFAD/*Neu1*^-/-^ and 5xFAD/GNIP had the highest beta-secretase activity, while female 5xFAD/CSFR and both genders of 5xFAD/NSE mice had the lowest activity (Extended Data Fig. 8b).

While there were similarities between alpha-and beta-secretase enzyme activity, gamma-secretase activity showed a distinct pattern. Compared to 5xFAD controls, female 5xFAD/CSFR and 5xFAD/NSE had lower levels of gamma-secretase activity, while 5xFAD/*Neu1*^-/-^ mice showed only a slightly reduction, and 5xFAD/GFAP and 5xFAD/GNIP had slightly higher levels (Fig. 8c). In males, 5xFAD/CSFR, 5xFAD/GFAP and 5xFAD/GNIP mice exhibited elevated gamma-secretase activity, while 5xFAD/NSE had reduced levels and 5xFAD/*Neu1*^-/-^ mice showed no significant change compared to 5xFAD controls (Fig. 8c). Gender comparisons within our genetic models revealed that female 5xFAD/*Neu1*^-/-^, 5xFAD/NSE, and 5xFAD/GFAP mice had higher gamma-secretase enzyme activity compared to males, while male 5xFAD/CSFR mice had higher enzyme activity than females, with no gender difference seen in 5xFAD/GNIP mice (Extended Data Fig. 8c). Among all 5xFAD models, 5xFAD/GNIP mice had the highest gamma-secretase activity, regardless of gender, while male 5xFAD/NSE mice had the lowest enzyme activity, followed closely by male 5xFAD/*Neu1*^-/-^ mice (Extended Data Fig. 8c).

Taken altogether, our findings highlight how cell-type specific alterations in PPCA and Neu1 expression differentially impact the enzyme activity levels of the secretases, while also revealing the inherent gender bias present. Additionally, we are now able to further explore the relationships observed between the levels of sialylated mature TACE, BACE1, and NCT, and their corresponding enzyme activities.

### Targeting NEU1 and PPCA via AAV-Mediated Gene Therapy to Reduce Plaque Burden in 5xFAD Mice

Our current findings, in combination with our previous study^38^, highlight that the overexpression of both NEU1 and PPCA, rather than either protein alone, offers the best therapeutic potential for mitigating AD pathology in 5xFAD mice. To further explore this therapeutic approach, we injected 3-month-old 5xFAD mice with an AAV2/8 bicistronic vector encoding both NEU1 and PPCA (5xFAD/AAV-PIN) and subsequently examined plaque pathology at 6-months of age. We observed robust expression of NEU1 and PPCA near the injection site, located close to the CA3 region of the hippocampus (Extended Data Fig. 8d). Overall, the total plaque area showed a trend towards reduction in 5xFAD/AAV-PIN mice, regardless of gender, but this reduction only reached significance in female mice (Fig. 8d and Extended Data Fig. 8d). Additionally, in female 5xFAD/AAV-PIN treated mice we observed a tendency toward a reduction in both the average plaque number per brain area unit and the average plaque size, while these measurements remained unchanged in male 5xFAD/AAV-PIN mice, compared to non-injected 5xFAD controls (Fig. 8e and 8f). No differences were observed in total brain area (Fig. 8g). One possible explanation for the more pronounced effect of this gene therapy approach in female mice, compared to males, is that females overall develop a higher pathologic burden, providing a more robust impact of AAV-PIN injection during a relatively short treatment period. Future studies aimed at assessing the long-term impacts of AAV-PIN injection in 5xFAD mice may more clearly demonstrate the therapeutic effectiveness of this approach, and solidify AAV-PIN gene therapy as a valuable therapeutic means for the treatment of both familial and sporadic AD.

## Discussion

AD remains an intractable neurodegenerative disorder with complex etiology and limited therapeutic options. Central to its pathology is the aberrant processing of APP, leading to the accumulation of amyloid peptides and subsequent plaque formation^1,10–12^. Despite extensive efforts to directly target Aβ production or aggregation, clinical translation has been largely disappointing^15,46^. Our study proposes a novel conceptual and mechanistic framework by implicating the interplay between Neu1 and PPCA, as key regulators of APP processing. These findings not only deepen our understanding of lysosomal involvement in AD but also uncover a potential therapeutic axis that connects glycosylation, enzyme activity, and neuroinflammation.

We demonstrate that the biochemical state of Neu1 and PPCA, defined by their expression, sialylation, and activity levels, plays a critical role in regulating the post-translational modification and turnover of APP and its secretases. In both human AD brain samples and 5xFAD/*Neu1^-/-^* mice, NEU1 deficiency is associated with increased sialylation of APP, BACE1, and TACE, which are key players in amyloidogenic and non-amyloidogenic cleavage pathways. This enhanced sialylation not only stabilizes these proteins but also potentiates their enzymatic activities, leading to accelerated amyloidogenic processing and increased production of Aβ42 peptides.

Importantly, we reveal that APP and its processing enzymes are not the only substrates of Neu1. PPCA itself is subject to Neu1-mediated de-sialylation, and its activity and turnover are similarly altered in Neu1-deficient contexts. This reciprocal dependency challenges the previously accepted unidirectional model, in which PPCA simply serves as a chaperone for Neu1. Our data instead suggest a tightly coordinated relationship, in which each protein stabilizes and regulates the other, forming a dynamic enzymatic partnership within the LMC. The state of this complex has profound implications not only for APP metabolism but also for overall lysosomal function.

One of the most compelling aspects of our study is the integration of glycosylation biology into the framework of AD pathogenesis. While previous research has demonstrated that glycosylation can affect protein trafficking and function^21–30^, the specific role of sialylation − a terminal, negatively charged glycosylation modification − has remained largely unexplored in the context of neurodegeneration. Here, we show that sialylation acts as a regulatory signal for APP cleavage and secretase activity. Increased sialylation enhances secretase enzymatic function, while de-sialylation through Neu1 dampens it. These findings are further supported by our data showing that Neu1 overexpression, or co-overexpression with PPCA, restores normal sialylation levels and attenuates the amyloidogenic cascade.

Additionally, we find that Neu1 dysregulation leads to a marked increase in lysosomal exocytosis, a mechanism through which lysosomal contents, including Aβ peptides and active enzymes, are secreted into the extracellular space^35,38,39^. In both AD patients and Neu1-deficient mice, we observed elevated levels of extracellular β-Hex activity, a marker of heightened lysosomal exocytosis activity. This process may contribute to neuroinflammation by releasing immunogenic substrates into the extracellular environment, thereby exacerbating microglial activation and neurotoxicity. Notably, recent studies propose that Aβ may function as a damage-associated molecular pattern (DAMP), capable of triggering autoimmune-like responses^47,48^. Our findings support this hypothesis by linking elevated lysosomal exocytosis to increased extracellular Aβ, suggesting that Neu1 and PPCA may serve as a critical node that connects proteostasis with immune signaling in AD.

Cell-type–specific overexpression of PPCA allowed us to further dissect the contributions of different brain cell populations to APP processing and Aβ dynamics. Neuron-specific expression (under the NSE promoter) was the most effective at reducing plaque burden, especially in female mice, underscoring the central role of neuronal APP metabolism in plaque pathology. In contrast, overexpression of PPCA in microglia or astrocytes altered enzyme sialylation and Aβ secretion but did not significantly affect plaque accumulation. These results point to a coordinated interplay between neuronal and glial cells in the regulation of LMC function and suggest that glial-targeted interventions may be more effective for modulating extracellular Aβ clearance or immune responses, rather than directly influencing primary APP processing.

Our data also underscore a clear and consistent gender disparity in NEU1/PPCA-related processes. Female mice exhibited higher baseline levels of sialylated APP and secretases, more pronounced lysosomal exocytosis activity, and greater plaque burden compared to male mice. These differences persisted across genetic backgrounds and experimental manipulations, suggesting a biologically intrinsic basis for the observed sex bias in AD prevalence and severity. While hormonal influences likely contribute, our findings underscore the importance of considering sex as a critical variable in therapeutic development and mechanistic studies.

In conclusion, our study positions NEU1 and PPCA, components of the LMC, as master regulators of APP processing, lysosomal function, and neuroinflammation in AD. By elucidating the reciprocal relationship between NEU1 and PPCA and revealing the critical role of protein sialylation in modulating secretase activity, we propose a paradigm shift in the conceptualization of amyloid pathology, from a cleavage-centric model to one that emphasizes enzymatic modulation through post-translational regulation. This new framework opens promising avenues for therapeutic development, particularly strategies aimed at restoring lysosomal homeostasis and fine-tuning glycosylation in the brain.

## Limitations

Despite the strengths of this study, several limitations warrant further investigation. First, β-galactosidase (β-Gal), a core member of the LMC, was not examined in this study. Its substrate, GM1 ganglioside, interacts with Aβ and may influence aggregation and toxicity.

Future work should assess how β-Gal expression and activity intersect with Neu1 and PPCA in the regulation of APP metabolism. Second, while our mouse models closely replicate human AD pathology, translation of these findings will require validation in human systems. In vitro studies using 3D Matrigel cultures or iPSC-derived organoids from AD patients could provide essential confirmation of Neu1/PPCA function and sialylation dynamics in a human context. Lastly, although our AAV-based approach proved effective in mice, optimization of vector delivery, distribution, and duration of expression will be essential for clinical translation.

## Supporting information

All supplemental figures with figure legends

## Acknowledgements

A.d’.A. holds the Jewelers for Children Endowed Chair in Genetics and Gene Therapy. This work was funded by the NIH grant 1RF1NS123174-01, The Assisi Foundation of Memphis, and the American Lebanese Syrian Associated Charities (ALSAC). The content is solely the responsibility of the authors and does not necessarily represent the official views of the National Institutes of Health. We thank Elida Gomero for maintaining the animals used in this study. We thank Dr. George Campbell, and the members of the Cell and Tissue Imaging Center–Light Microscopy Shared Resource of SJCRH for their assistance with processing and imaging tissue. We are grateful to Dr. Randall Woltjer and the Oregon Brain Bank at Oregon Health and Science University, and to the Alzheimer’s Disease Research Center at University of California Davis.

## Author Contributions

Conceptualization, L.E.F. I.A. and A.d’A. Methodology, L.E.F., D.v.d.V., H.H., J.A.W., I.A., and A.d’A. Validation, L.E.F. D.v.d.V. G.Y. and A.d’A. Formal Analysis, L.E.F. D.v.d.V. and J.A.W. Investigation, L.E.F., D.v.d.V., I.A., and A.d’A. Resources, L.E.F., D.v.d.V., H.H., J.A.W. G.Y. and A.d’A. Data Curation, L.E.F. D.v.d.V., and A.d’A. Writing – Original Draft Preparation, L.E.F. Writing – Review & Editing, L.E.F. D.v.d.V., G.Y. and A.d’A. Visualization, L.E.F., D.v.d.V., and A.d’A. Supervision, A.d’A. Project Administration, A.d’A. Funding Acquisition, A.d’A. All authors have read and agreed to the published version of the manuscript.

## Materials and Methods

### Animal Models

Animals were housed in a fully AAALAC (Assessment and Accreditation of Laboratory Animal Care)-accredited animal facility with controlled temperature (22°C), humidity, and lighting (alternating 12-h light/dark cycles). Food and water were provided *ad libitum*. All procedures in mice were performed according to animal protocols approved by the St.

Jude Children’s Research Hospital Institutional Animal Care and Use Committee and NIH guidelines. Sexes were represented equally and characterized separately. WT and *Neu1^-/-^* mice were bred into the C57BL/6J background. CSFR^PPCA-TG^, GFAP^PPCA-TG^, NSE^PPCA-TG^, and GNIP^Neu^^1-PPCA-TG^ were bred into the FVB background. 5xFAD mice were obtained from Jackson Laboratories and bred into the SJL-C57BL background. *Neu1^-/-^*, CSFR^PPCA-TG^, GFAP^PPCA-TG^, NSE^PPCA-TG^, and GNIP^Neu^^1-PPCA-TG^ were crossed with 5xFAD mice to produce the crosses analyzed in this manuscript. Separate WT and 5xFAD controls were collected between the Neu1^-/-^ and PPCA overexpression lines to account for differences between the C57BL/6J versus FVB backgrounds.

### Immunohistochemical Analyses

Mouse brains were collected and kept in 10% formalin until paraffin embedding. Immunohistochemical (IHC) analyses were performed on 6-μm-thick serial paraffin sections. Sections were subjected to automated de-paraffinization and antigen retrieval using a 20-min cycle in a pressure cooker with slides submerged in 10 mM sodium citrate/0.1% Tween-20. After washing sections 3 times for 5 min in PBS, they were blocked in Horse Serum for 1 h at RT then incubated overnight at RT with antibodies against amyloid (4G8, 1:1000), human LAMP1 (1:300), human NEU1 (1:100), or human PPCA (1:200) diluted in blocking buffer. The sections were then washed 3 times for 5 min with PBS and incubated with biotinylated secondary antibody for 2 h at RT. Endogenous peroxidase was quenched by incubating the sections with 5% hydrogen peroxidase/PBS for 15 min at RT. Antibodies were detected using the VECTASTAIN^®^ Elite^®^ ABC Kit, and diaminobenzidine substrate, and sections were counterstained with hematoxylin per standard methods.

### Enzyme Activity Assays

CSF from 6-month-old were collected and used for enzymatic assays of β-hex activity. CSF (10 μL) was incubated with 10 μL 4MU-N-acetyl-β-D-glucosaminide for 1 h at 37C. To stop enzyme reactions, 200 μL 0.5 M carbonate buffer (pH 10.7) was added to all wells. Fluorescence was measured on a plate reader (EX-355 nm, EM-460 nm). The net fluorescence values were compared with those of the linear 4MU standard curve and were used to calculate the specific enzyme activities. Activities were calculated as nanomoles of substrate converted per hour per milligram of protein.

Hippocampal tissue was lysed in ddH_2_O for Neu1, PPCA, TACE, and BACE1 activity assays. Neu1 activity was measured by incubating 5uL of lysates with 5 μL 4MU-α-D-N-acetylneuraminic acid at pH 4.3, To stop enzyme reactions, 200 μL 0.5 M carbonate buffer (pH 10.7) was added to all wells. Fluorescence was measured on a plate reader (EX-355 nm, EM-460 nm). The net fluorescence values were compared with those of the linear 4MU standard curve and were used to calculate the specific enzyme activities. Activities were calculated as nanomoles of substrate converted per hour per milligram of protein.

PPCA activity was measured by incubating 10μL of lysates with 10μL Z-Phe-Ala. Enzyme reactions were stopped at 100°C for 5 min and 10 μL reaction mixture was read in 250 μL 50 mM Na-carbonate stop buffer (pH 9.5) containing 500 μL *o*-phtaldialdehyde (10 mg/mL) and 500 μL 2-mercaptoethanol (5 μL/mL) per 30 mL. Fluorescence was measured on a plate reader (EX-355 nm, EM-460 nm). The net fluorescence values were compared with those of the linear standard curve and were used to calculate the specific enzyme activities. Activities were calculated as nanomoles of substrate converted per hour per milligram of protein.

TACE and BACE1 activity were measured with commercially available kits, BML-AK310 and AS-71144, respectively. Briefly, 20μL of lysate was added to 30μL of pre-warmed assay buffer, then 50μL of diluted substrate solution was added to each well and plates were incubated at 37C for 3 h (TACE) or overnight (BACE1). Fluorescence was measured on a plate reader, TACE at EX-328 nm/EM-420 nm and BACE1 at EX-490 nm/EM-520 nm. The net fluorescence values were compared with those of the linear standard curve and were used to calculate the specific enzyme activities. Activities were calculated as nanomoles of substrate converted per hour per milligram of protein.

Hippocampal tissue was lysed in 1x CHAPS buffer for gamma-secretase activity assays. Activity was measured by incubating 20μL of lysate with 20μL of NMA-GGVVIATVK(DNP)-*D*R *D*R *D*R-NH₂ in CHAPS buffer overnight at 37C. Fluorescence was measured on a plate reader (EX-355 nm, EM-440 nm). The net fluorescence values were compared with those of the linear standard curve and were used to calculate the specific enzyme activities. Activities were calculated as nanomoles of substrate converted per hour per milligram of protein.

### CSF collection

For collection of CSF, mice were anaesthetized with avertin and placed into a Kopf stereotaxic device. A small incision was made at the base of the skull, and CSF was collected from the cisterna magna by using a syringe attached to a butterfly needle. Collected CSF was dispensed into 250 μL Eppendorf tubes, frozen in liquid nitrogen, and then stored at –80°C until needed.

### Cycloheximide Treatments of Hippocampal Slice Cultures

Hippocampi were collected into collection media and sectioned into 300 μm sections then placed on culture insets in pre-warmed media in 6-well culture dishes. Hippocampal slice cultures were left for two weeks before proceeding.

For CHX assays, media was replaced with new media containing 100 μM CHX and incubated at 37°C in 5% CO_2_ for 4, 8, or 16 h; untreated hippocampal slices were used as 0 h controls. At the appropriate time point, slices were washed with PBS and collected into RIPA lysis buffer containing protease inhibitors. Protein concentrations were determined using bicinchoninic acid (BCA) assay, then samples containing 30 μg total protein were prepared for immunoblot analysis.

### Enzyme-linked Immunosorbent Assays

Aβ42 concentrations were measured in the CSF collected from 6-month-old mice using a commercially available kit (KMB3441). Aβ42 levels in tested CSF samples were determined using the sandwich enzyme-linked immunosorbent assay (ELISA) kits per the manufacturer’s instructions. Briefly, 100 μL/well of diluted sample or assay standard was added to 96-well plates and incubated for 2 h at RT. Plates were washed 4 times with 400 μL wash buffer and then incubated for 1 h with 100 μL/well of the detection antibody solution at RT. Plates were rinsed 4 times with wash buffer and incubated with 100 μL/well of the HRP solution at RT. Plates were rinsed 4 times with wash buffer and incubated with 100 μL/well of the stabilized chromogen solution at RT in the dark. After this period, 100 μL/well of stop solution was added to terminate the reaction. The absorbance was measured at 450 nm.

### Immunoblotting

Hippocampi from 6-month-old mice were homogenized with RIPA buffer containing protease inhibitors. Protein concentrations were determined using a BCA assay. Proteins were separated by SDS–PAGE on precast Criterion Midi 4%-20% or 10% gels under reducing conditions and transferred to a polyvinylidene difluoride membrane.

Membranes were incubated for 1 h in blocking buffer (5% dry milk in TBS-Tween) at RT and subsequently probed with antibodies against huPPCA (1:200), moPPCA (1:10), APP (1:500), TACE (1:500), BACE1 (1:500), and NCT (1:500) diluted in 5% BSA/TBS-Tween overnight at 4°C. Following this step, blots were washed 3 times for 5 min with TBS-Tween and then incubated with the appropriate secondary antibody for 2 h at RT. Immunoblots were developed using the Clarity Max^TM^ Western ECL substrate.

### Immunoprecipitation of Sialylated Proteins

6-month-old hippocampi were homogenized with RIPA buffer containing protease inhibitors. Protein concentrations were determined using a BCA assay. For each SNA immunoprecipitation, 200 μg total lysate was used. Samples were rotated at 4°C for 16 h with 2 μg of biotinylated SNA, in a final volume of 500 mL RIPA + 1% BSA. After incubation, 10 μL washed Nanolink™ streptavidin magnetic beads, diluted (1:2) in RIPA, was added to each sample. Samples were rotated for 3 h at RT and then placed on a magnet. Beads were washed 4 times in 0.1x RIPA and then resuspended in 20 μL LDS running buffer and boiled for 5 min at 95°C. Samples where then spun down, placed on a magnet, and loaded on SDS-PAGE gels to be analyzed by immunoblot; samples were prepared accordingly, containing 15 μg total protein for input controls.

### Vector Production

The *scAAV2/8-CMV*-*CTSA-IRES-NEU1* (huPPCA and huNEU1) construct contains a cytomegalovirus (CMV) promoter, which ensured the expression of the 1.44-kb human *PPCA* complementary DNA and the 1.247-kb human *NEU1* cDNA. The scAAV vector particles were made in the Children’s GMP, LLC facility on the St. Jude campus. The genome titer of the vector (5.69 × 10^12^ GC/mL) was determined by qPCR and/or direct loading and electrophoresis of detergent-treated vector particles on native agarose gels, staining with fluorescent dye, and quantification of signal relative to known mass standards.

### AAV Transduction *In Vivo*

Animals were anaesthetized and placed into a Kopf stereotaxic device. Intracerebroventricular injections were done unilaterally into the left hippocampus. Injections were performed with a 10-μL Hamilton syringe fitted with a glass micropipette. 3 × 10^10^ GC of virus diluted in 10 μL of sterile PBS was injected per mouse. After each injection, the needle was left in place for 5 min before withdrawal from the brain. Mice were housed individually for 3 months after the injection, then anesthetized, perfused with ice-cold PBS, and their brains were processed for downstream analyses.

### Quantification and Statistical Analyses

Statistical analyses were performed using GraphPad Prism 10 software. Quantitative data are represented as mean ± SD. For comparisons between control and test, Welch’s student’s *t*-tests (unpaired, 2-tailed) were performed. For comparisons of controls and multiple tests within one sex, 1-way ANOVA with multiple comparisons was performed. For comparisons of sexes between multiple controls and tests, 2-way ANOVA with multiple comparisons was performed. For comparisons of sexes between multiple controls and tests, where time was a third variable, 3-way ANOVA was performed. P-values <0.05 were considered statistically significant. The number of replicates for a given experiment is indicated in each Fig. legend.

## References

1 Muller, U. C. & Zheng, H. Physiological functions of APP family proteins. Cold Spring Harb Perspect Med 2, a006288 (2012). 10.1101/cshperspect.a006288

2 Turner, P. R., O’Connor, K., Tate, W. P. & Abraham, W. C. Roles of amyloid precursor protein and its fragments in regulating neural activity, plasticity and memory. Prog Neurobiol 70, 1–32 (2003). 10.1016/s0301-0082(03)00089-3

3 Priller, C. et al. Synapse formation and function is modulated by the amyloid precursor protein. J Neurosci 26, 7212–7221 (2006). 10.1523/JNEUROSCI.1450-06.2006

4 Satpute-Krishnan, P., DeGiorgis, J. A., Conley, M. P., Jang, M. & Bearer, E. L. A peptide zipcode sufficient for anterograde transport within amyloid precursor protein. Proc Natl Acad Sci U S A 103, 16532–16537 (2006). 10.1073/pnas.0607527103

5 Duce, J. A. et al. Iron-export ferroxidase activity of beta-amyloid precursor protein is inhibited by zinc in Alzheimer’s disease. Cell 142, 857–867 (2010). 10.1016/j.cell.2010.08.014

6 Zhang, T., Chen, D. & Lee, T. H. Phosphorylation Signaling in APP Processing in Alzheimer’s Disease. Int J Mol Sci 21 (2019). 10.3390/ijms21010209

7 Ohline, S. M. et al. Effect of soluble amyloid precursor protein-alpha on adult hippocampal neurogenesis in a mouse model of Alzheimer’s disease. Mol Brain 15, 5 (2022). 10.1186/s13041-021-00889-1

8 Cha, H. J., Shen, J. & Kang, J. Regulation of gene expression by the APP family in the adult cerebral cortex. Sci Rep 12, 66 (2022). 10.1038/s41598-021-04027-8

9 Kuhn, A. J., Abrams, B. S., Knowlton, S. & Raskatov, J. A. Alzheimer’s Disease “Non-amyloidogenic” p3 Peptide Revisited: A Case for Amyloid-alpha. ACS Chem Neurosci 11, 1539–1544 (2020). 10.1021/acschemneuro.0c00160

10 Im, E. et al. Lysosomal dysfunction in Down syndrome and Alzheimer mouse models is caused by v-ATPase inhibition by Tyr(682)-phosphorylated APP betaCTF. Sci Adv 9, eadg1925 (2023). 10.1126/sciadv.adg1925

11 Lee, S. E. et al. Accumulation of APP-CTF induces mitophagy dysfunction in the iNSCs model of Alzheimer’s disease. Cell Death Discov 8, 1 (2022). 10.1038/s41420-021-00796-3

12 Chen, G. F. et al. Amyloid beta: structure, biology and structure-based therapeutic development. Acta Pharmacol Sin 38, 1205–1235 (2017). 10.1038/aps.2017.28

13 Prosswimmer, T., Heng, A. & Daggett, V. Mechanistic insights into the role of amyloid-beta in innate immunity. Sci Rep 14, 5376 (2024). 10.1038/s41598-024-55423-9

14 Behl, C. In 2024, the amyloid-cascade-hypothesis still remains a working hypothesis, no less but certainly no more. Front Aging Neurosci 16, 1459224 (2024). 10.3389/fnagi.2024.1459224

15 Zhang, J. et al. Recent advances in Alzheimer’s disease: Mechanisms, clinical trials and new drug development strategies. Signal Transduct Target Ther 9, 211 (2024). 10.1038/s41392-024-01911-3

16 Yang, K. F. et al. Secretase promotes AD progression: simultaneously cleave Notch and APP. Front Aging Neurosci 16, 1445470 (2024). 10.3389/fnagi.2024.1445470

17 Shah, S. et al. Nicastrin functions as a gamma-secretase-substrate receptor. Cell 122, 435–447 (2005). 10.1016/j.cell.2005.05.022

18 Lu, P. et al. Three-dimensional structure of human gamma-secretase. Nature 512, 166–170 (2014). 10.1038/nature13567

19 Zhu, W., Zhou, Y., Guo, L. & Feng, S. Biological function of sialic acid and sialylation in human health and disease. Cell Death Discov 10, 415 (2024). 10.1038/s41420-024-02180-3

20 Yang, K., Yang, Z., Chen, X. & Li, W. The significance of sialylation on the pathogenesis of Alzheimer’s disease. Brain Res Bull 173, 116–123 (2021). 10.1016/j.brainresbull.2021.05.009

21 Haukedal, H. & Freude, K. K. Implications of Glycosylation in Alzheimer’s Disease. Front Neurosci 14, 625348 (2020). 10.3389/fnins.2020.625348

22 Fastenau, C. et al. Distinct patterns of plaque and microglia glycosylation in Alzheimer’s disease. Brain Pathol 34, e13267 (2024). 10.1111/bpa.13267

23 Nakagawa, K. et al. Sialylation enhances the secretion of neurotoxic amyloid-beta peptides. J Neurochem 96, 924–933 (2006). 10.1111/j.1471-4159.2005.03595.x

24 Vanoni, O., Paganetti, P. & Molinari, M. Consequences of individual N-glycan deletions and of proteasomal inhibition on secretion of active BACE. Mol Biol Cell 19, 4086–4098 (2008). 10.1091/mbc.e08-05-0459

25 Charlwood, J. et al. Characterization of the glycosylation profiles of Alzheimer’s beta-secretase protein Asp-2 expressed in a variety of cell lines. J Biol Chem 276, 16739–16748 (2001). 10.1074/jbc.M009361200

26 Chavaroche, A. et al. Glycosylation of a disintegrin and metalloprotease 17 affects its activity and inhibition. Anal Biochem 449, 68–75 (2014). 10.1016/j.ab.2013.12.018

27 Xie, T. et al. Crystal structure of the gamma-secretase component nicastrin. Proc Natl Acad Sci U S A 111, 13349–13354 (2014). 10.1073/pnas.1414837111

28 Herreman, A. et al. gamma-Secretase activity requires the presenilin-dependent trafficking of nicastrin through the Golgi apparatus but not its complex glycosylation. J Cell Sci 116, 1127–1136 (2003). 10.1242/jcs.00292

29 Moniruzzaman, M., Ishihara, S., Nobuhara, M., Higashide, H. & Funamoto, S. Glycosylation status of nicastrin influences catalytic activity and substrate preference of gamma-secretase. Biochem Biophys Res Commun 502, 98–103 (2018). 10.1016/j.bbrc.2018.05.126

30 Zhao, J. & Lang, M. New insight into protein glycosylation in the development of Alzheimer’s disease. Cell Death Discov 9, 314 (2023). 10.1038/s41420-023-01617-5

31 Marques, A. R. A. & Saftig, P. Lysosomal storage disorders - challenges, concepts and avenues for therapy: beyond rare diseases. J Cell Sci 132 (2019). 10.1242/jcs.221739

32 Platt, F. M., d’Azzo, A., Davidson, B. L., Neufeld, E. F. & Tifft, C. J. Lysosomal storage diseases. Nat Rev Dis Primers 4, 27 (2018). 10.1038/s41572-018-0025-4

33 de Geest, N. et al. Systemic and neurologic abnormalities distinguish the lysosomal disorders sialidosis and galactosialidosis in mice. Hum Mol Genet 11, 1455–1464 (2002). 10.1093/hmg/11.12.1455

34 Bonten, E. J. et al. Heterodimerization of the sialidase NEU1 with the chaperone protective protein/cathepsin A prevents its premature oligomerization. J Biol Chem 284, 28430–28441 (2009). 10.1074/jbc.M109.031419

35 Yogalingam, G. et al. Neuraminidase 1 is a negative regulator of lysosomal exocytosis. Dev Cell 15, 74–86 (2008). 10.1016/j.devcel.2008.05.005

36 Cuervo, A. M., Mann, L., Bonten, E. J., d’Azzo, A. & Dice, J. F. Cathepsin A regulates chaperone-mediated autophagy through cleavage of the lysosomal receptor. EMBO J 22, 47–59 (2003). 10.1093/emboj/cdg002

37 Wu, J., Xu, W., Su, Y., Wang, G. H. & Ma, J. J. Targeting chaperone-mediated autophagy in neurodegenerative diseases: mechanisms and therapeutic potential. Acta Pharmacol Sin (2024). 10.1038/s41401-024-01416-3

38 Annunziata, I. et al. Lysosomal NEU1 deficiency affects amyloid precursor protein levels and amyloid-beta secretion via deregulated lysosomal exocytosis. Nat Commun 4, 2734 (2013). 10.1038/ncomms3734

39 Fremuth, L. E. et al. Neuraminidase 1 regulates neuropathogenesis by governing the cellular state of microglia via modulation of Trem2 sialylation. Cell Rep 44, 115204 (2025). 10.1016/j.celrep.2024.115204

40 Tachida, Y. et al. O-GalNAc glycosylation determines intracellular trafficking of APP and Abeta production. J Biol Chem 299, 104905 (2023). 10.1016/j.jbc.2023.104905

41 Shi, J. et al. Comprehensive analysis of O-glycosylation of amyloid precursor protein (APP) using targeted and multi-fragmentation MS strategy. Biochim Biophys Acta Gen Subj 1865, 129954 (2021). 10.1016/j.bbagen.2021.129954

42 Wang, D., Zaitsev, S., Taylor, G., d’Azzo, A. & Bonten, E. Protective protein/cathepsin A rescues N-glycosylation defects in neuraminidase-1. Biochim Biophys Acta 1790, 275–282 (2009). 10.1016/j.bbagen.2009.01.006

43 Bonten, E. J., Annunziata, I. & d’Azzo, A. Lysosomal multienzyme complex: pros and cons of working together. Cell Mol Life Sci 71, 2017–2032 (2014). 10.1007/s00018-013-1538-3

44 Hahn, C. N., del Pilar Martin, M., Zhou, X. Y., Mann, L. W. & d’Azzo, A. Correction of murine galactosialidosis by bone marrow-derived macrophages overexpressing human protective protein/cathepsin A under control of the colony-stimulating factor-1 receptor promoter. Proceedings of the National Academy of Sciences of the United States of America 95, 14880–14885 (1998). 10.1073/pnas.95.25.14880

45 Su, M. et al. Expression specificity of GFAP transgenes. Neurochem Res 29, 2075–2093 (2004). 10.1007/s11064-004-6881-1

46 Yiannopoulou, K. G. & Papageorgiou, S. G. Current and Future Treatments in Alzheimer Disease: An Update. J Cent Nerv Syst Dis 12, 1179573520907397 (2020). 10.1177/1179573520907397

47 Huang, Y. M. et al. Amyloids in Site-Specific Autoimmune Reactions and Inflammatory Responses. Front Immunol 10, 2980 (2019). 10.3389/fimmu.2019.02980

48 Weaver, D. F. beta-Amyloid is an Immunopeptide and Alzheimer’s is an Autoimmune Disease. Curr Alzheimer Res 18, 849–857 (2021). 10.2174/1567205018666211202141650

