## Supplementary material for "Targeting Lysosomal Dysfunction to Alleviate Plaque Deposition in an Alzheimer Disease Model": All supplemental figures with figure legends

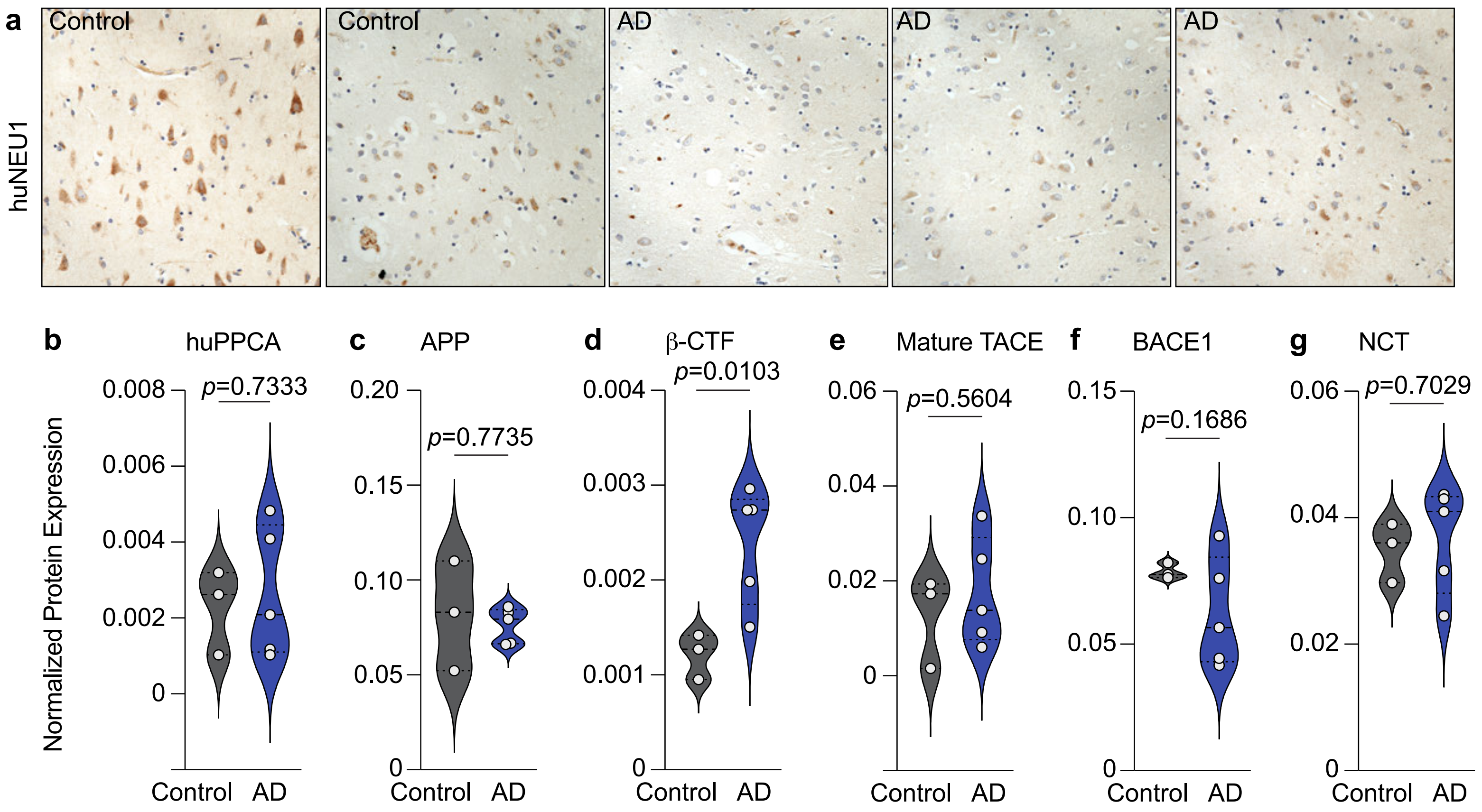
a
Control
Control
AD
AD
AD
huNEU1
b
huPPCA
p=0.7333
c
APP
p=0.7735
d
β-CTF
p=0.0103
e
Mature TACE
p=0.5604
f
BACE1
p=0.1686
g
NCT
p=0.7029
Normalized Protein Expression
Control
AD

**Extended Data Fig. 1: Total levels of AD-associated proteins in human AD patient brains.**

**a**, Immunohistochemical staining of NEU1 (brown) in human controls (n=10) and AD patients (n=10). **b-g**, Measurements collected from in homogenized hippocampal tissue of controls (n=3) and AD patients (n=5). Total normalized protein expression **b**, huPPCA. **c**, APP. **d**, β-CTF. **e**, mature TACE. **f**, BACE1. **g**, NCT. Actual p-values are given, any value less than or equal to 0.05 is considered significant.

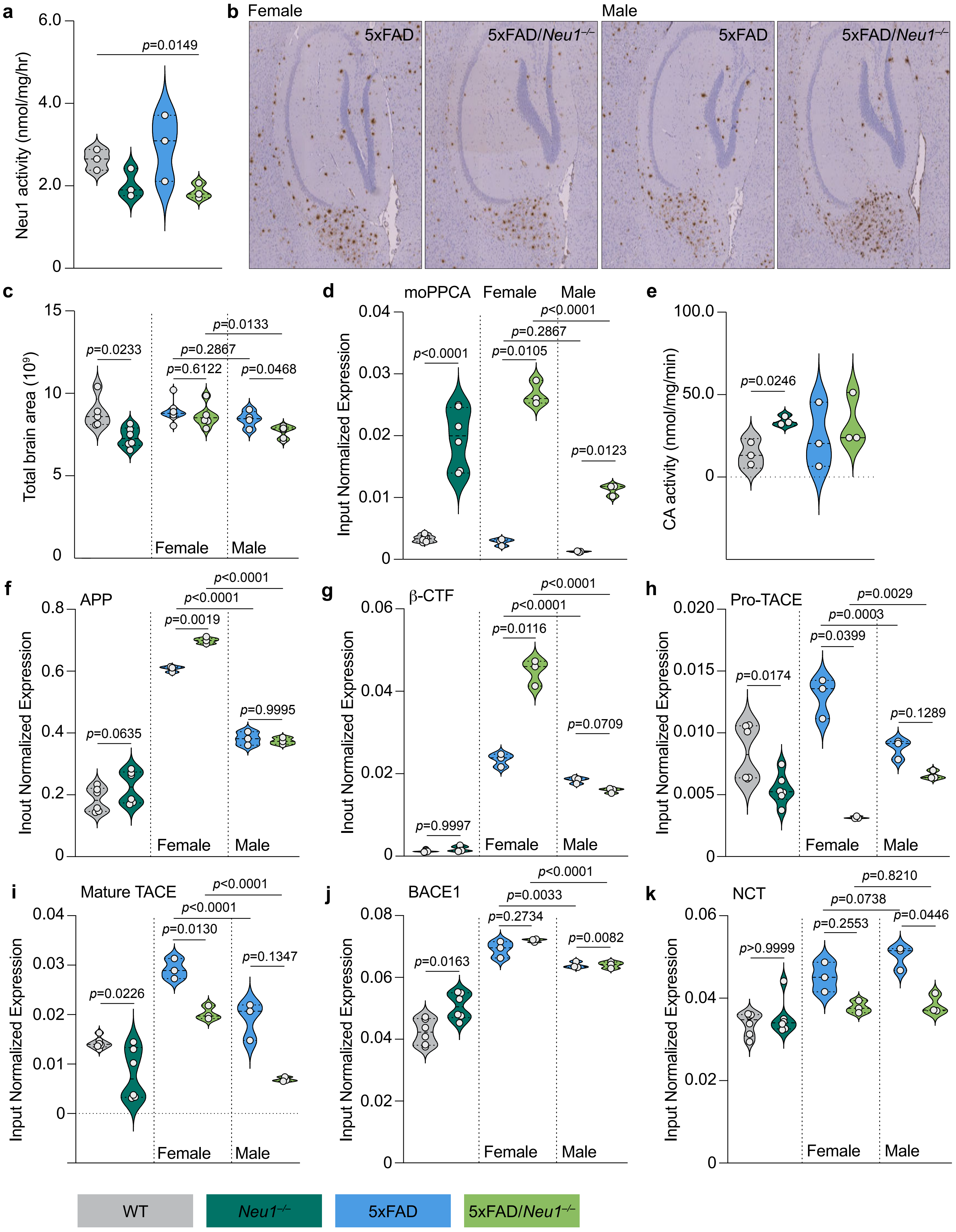
a
Neu1 activity (nmol/mg/hr)
p=0.0149
b Female
Male
5xFAD
5xFAD/Neu1−/−
c
Total brain area (10⁹)
p=0.0233
p=0.6122
p=0.2867
p=0.0133
p=0.0468
Female
Male
d
moPPCA
Input Normalized Expression
p<0.0001
p=0.0105
p=0.2867
p<0.0001
p=0.0123
e
CA activity (nmol/mg/min)
p=0.0246
f
APP
Inout Normalized Expression
p=0.0635
p=0.0019
p<0.0001
p<0.0001
p=0.9995
g
β-CTF
p=0.9997
p=0.0116
p<0.0001
p<0.0001
p=0.0709
h
Pro-TACE
p=0.0174
p=0.0399
p=0.0003
p=0.0029
p=0.1289
i
Mature TACE
p=0.0226
p=0.0130
p<0.0001
p<0.0001
p=0.1347
j
BACE1
p=0.0163
p=0.2734
p=0.0033
p<0.0001
p=0.0082
k
NCT
p>0.9999
p=0.2553
p=0.0738
p=0.8210
p=0.0446
WT
Neu1−/−
5xFAD
5xFAD/Neu1−/−

**Extended Data Fig. 2: Analysis of total proteins in 5xFAD mice when Neu1 is deficient.**

**a**, Quantification of Neu1 enzyme activity in homogenized hippocampal lysates from WT, *Neu1*$^{-/-}$, 5xFAD, and 5xFAD/*Neu1*$^{-/-}$ mice (n=3 for all). **b** and **c**, Collected from paraffin-embedded whole brain sections for WT (n=5), Neu1$^{-/-}$ (n=6), female 5xFAD (n=8), female 5xFAD/*Neu1*$^{-/-}$ (n=6), male 5xFAD (n=4) and male 5xFAD/*Neu1*$^{-/-}$ (n=6) mice. **b**, Immunohistochemical staining of amyloid (4G8; brown). **c**, Quantification of total brain area. **d**, Total normalized protein expression of PPCA in homogenized hippocampal lysates from WT (n=6), Neu1$^{-/-}$ (n=6), female 5xFAD (n=3), female 5xFAD/*Neu1*$^{-/-}$ (n=3), male 5xFAD (n=3) and male 5xFAD/*Neu1*$^{-/-}$ (n=3) mice. **e**, Quantification of PPCA (Cathepsin A) enzyme activity in homogenized hippocampal lysates from WT, *Neu1*$^{-/-}$, 5xFAD, and 5xFAD/*Neu1*$^{-/-}$ mice (n=3 for all). **f-k**, Measurements collected from homogenized hippocampal lysates from WT (n=6), *Neu1*$^{-/-}$ (n=6), female 5xFAD (n=3), female 5xFAD/*Neu1*$^{-/-}$ (n=3), male 5xFAD (n=3) and male 5xFAD/*Neu1*$^{-/-}$ (n=3) mice. Total normalized protein expression **f**, APP. **g**, $\beta$-CTF. **h**, pro-TACE, **i**, mature TACE. **j**, BACE1. **k**, NCT. Actual p-values are given, any value less than or equal to 0.05 is considered significant.

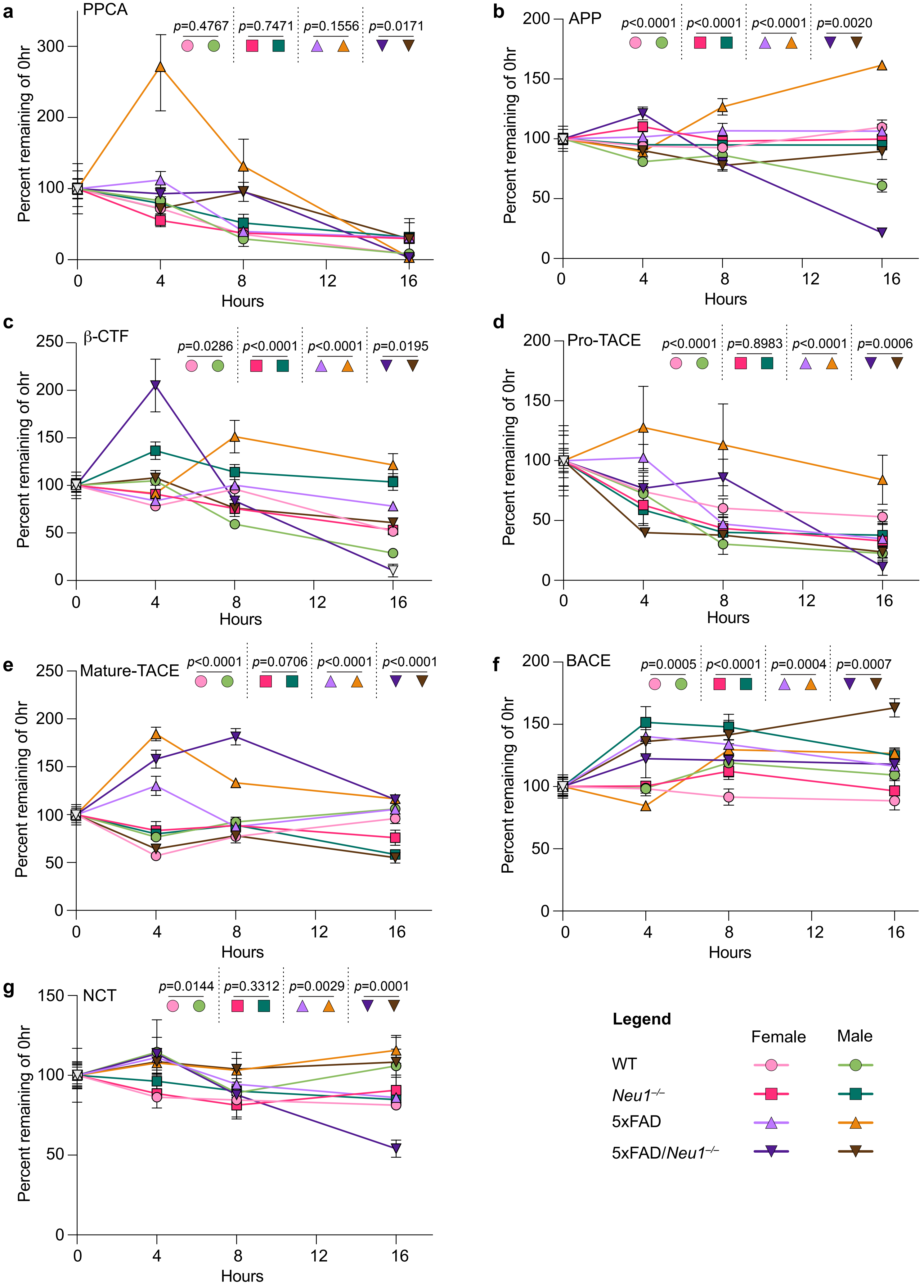
a
PPCA
p=0.4767
p=0.7471
p=0.1556
p=0.0171
Percent remaining of 0hr
Hours
b
APP
p<0.0001
p<0.0001
p<0.0001
p=0.0020
Percent remaining of 0hr
Hours
c
β-CTF
p=0.0286
p<0.0001
p<0.0001
p=0.0195
Percent remaining of ohr
Hours
d
Pro-TACE
p<0.0001
p=0.8983
p<0.0001
p=0.0006
Percent remaining of 0hr
Hours
e
Mature-TACE
p<0.0001
p=0.0706
p<0.0001
p<0.0001
Percent remaining of 0hr
Hours
f
BACE
p=0.0005
p<0.0001
p=0.0004
p=0.0007
Percent remaining of 0hr
Hours
g
NCT
p=0.0144
p=0.3312
p=0.0029
p=0.0001
Percent remaining of 0hr
Hours
Legend
Female
Male
WT
Neu1−/−
5xFAD
5xFAD/Neu1−/−

**Extended Data Fig. 3: Comparison of protein turnover between females and males when *Neu1* is deficient in 5xFAD mice.**

**a-g**, Measurements collected from hippocampal slice cultures of female WT (n=3), female *Neu1*$^{-/-}$ (n=3), female 5xFAD (n=3), female 5xFAD/*Neu1*$^{-/-}$ (n=3), male WT (n=3), male *Neu1*$^{-/-}$ (n=3), male 5xFAD (n=3) and male 5xFAD/*Neu1*$^{-/-}$ (n=3) mice treated with 100μM cycloheximide (CHX) for 0, 4, 8, and 16 h to calculate protein turnover, displayed as a percentage of quantified protein remaining compared to the 0 h timepoint. Protein turnover plots for **a**, moPPCA. **b**, APP. **c**, β-CTF. **d**, pro-TACE. **e**, mature TACE. **f**, BACE1. **g**, NCT. Female and male time course plots have been combined for comparison between sexes rather than genetic background. p-values represent significance of male to female statistical comparisons.

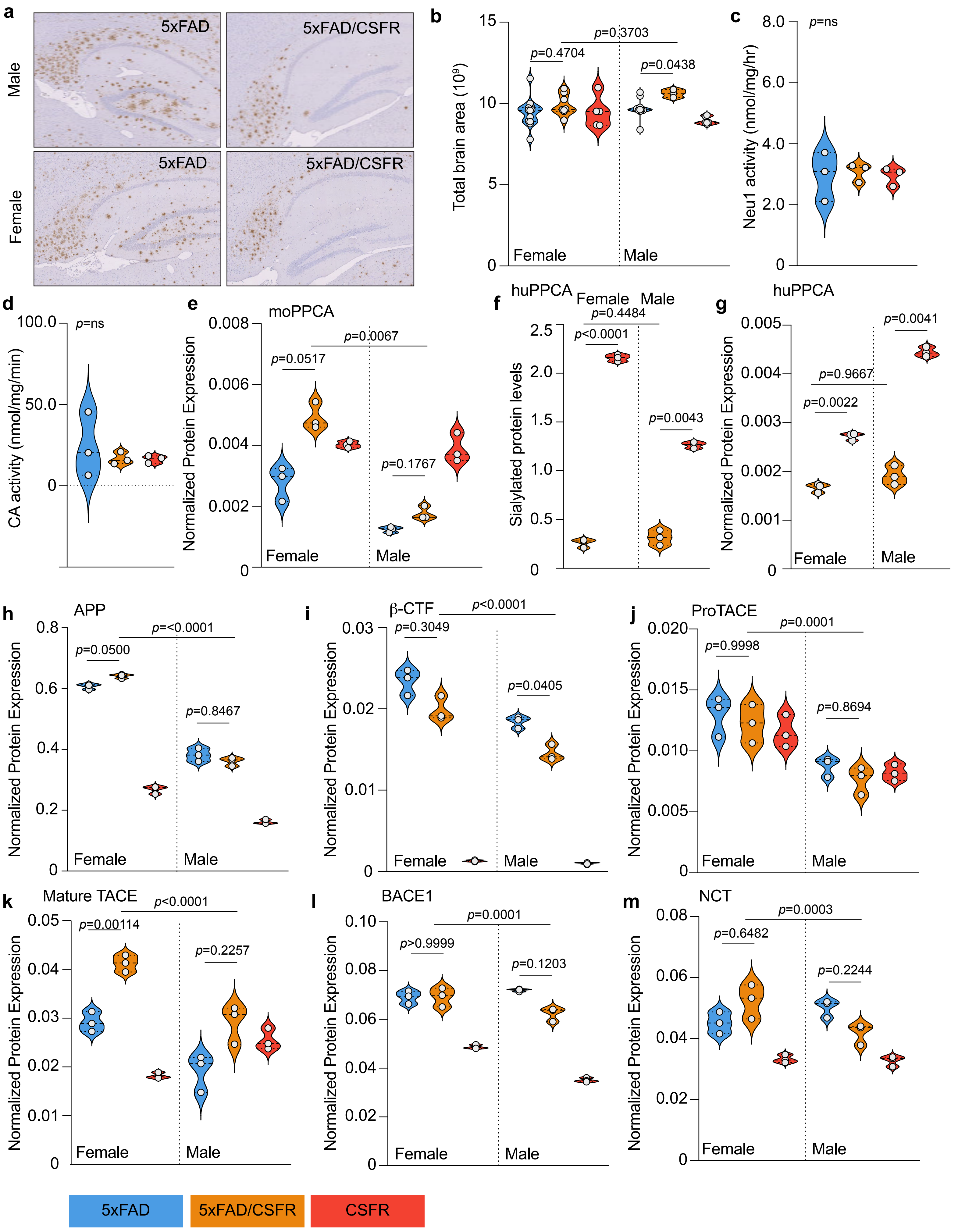

a
Male
Female
5xFAD
5xFAD/CSFR
b
Total brain area (10⁹)
p=0.4704
p=0.3703
p=0.0438
Female
Male
c
Neu1 activity (nmol/mg/hr)
p=ns
d
CA activity (nmol/mg/min)
p=ns
e
moPPCA
Normalized Protein Expression
p=0.0517
p=0.0067
p=0.1767
f
huPPCA
Sialylated protein levels
p=0.4484
p<0.0001
p=0.0043
g
huPPCA
p=0.0041
p=0.9667
p=0.0022
h
APP
p=0.0500
p=<0.0001
p=0.8467
i
β-CTF
p<0.0001
p=0.3049
p=0.0405
j
ProTACE
p=0.0001
p=0.9998
p=0.8694
k
Mature TACE
p<0.0001
p=0.00114
p=0.2257
l
BACE1
p=0.0001
p>0.9999
p=0.1203
m
NCT
p=0.0003
p=0.6482
p=0.2244
5xFAD
5xFAD/CSFR
CSFR

**Extended Data Fig. 4: Analysis of total proteins in 5xFAD mice when PPCA is overexpressed under the CSFR1 promoter.**

**a** and **b**, Collected from paraffin-embedded whole brain sections for female 5xFAD (n=12), female 5xFAD/CSFR (n=8), female CSFR (n=5), male 5xFAD (n=10), male 5xFAD/CSFR (n=3), and male CSFR (n=6) mice. **a**, Immunohistochemical staining of amyloid (4G8; brown). **b**, Quantification of total brain area. **c** and **d**, Measurements collected from homogenized hippocampal lysates of 5xFAD, 5xFAD/CSFR, and CSFR mice (n=3 for all). **c**, Quantification of Neu1 enzyme activity. **d**, Quantification of PPCA (Cathepsin A) enzyme activity. **e-m**, Measurements collected from homogenized hippocampal lysates of female 5xFAD (n=3), female 5xFAD/CSFR (n=3), female CSFR (n=3), male 5xFAD (n=3), male 5xFAD/CSFR (n=3), and male CSFR (n=3) mice. **e**, Total normalized protein expression of moPPCA. **f**, Normalized expression of sialylated huPPCA, relative to total protein levels displayed in *Extended Data Fig. 4g*. 5xFAD does not express huPPCA so no data was shown for this group. **g**, Total normalized protein expression of huPPCA. 5xFAD does not express huPPCA so no data was shown for this group. **h-m**, Total normalized protein expression **h**, APP. **i**, β-CTF. **j**, pro-TACE, **k**, mature TACE. **l**, BACE1. **m**, NCT. Actual p-values are given, any value less than or equal to 0.05 is considered significant.

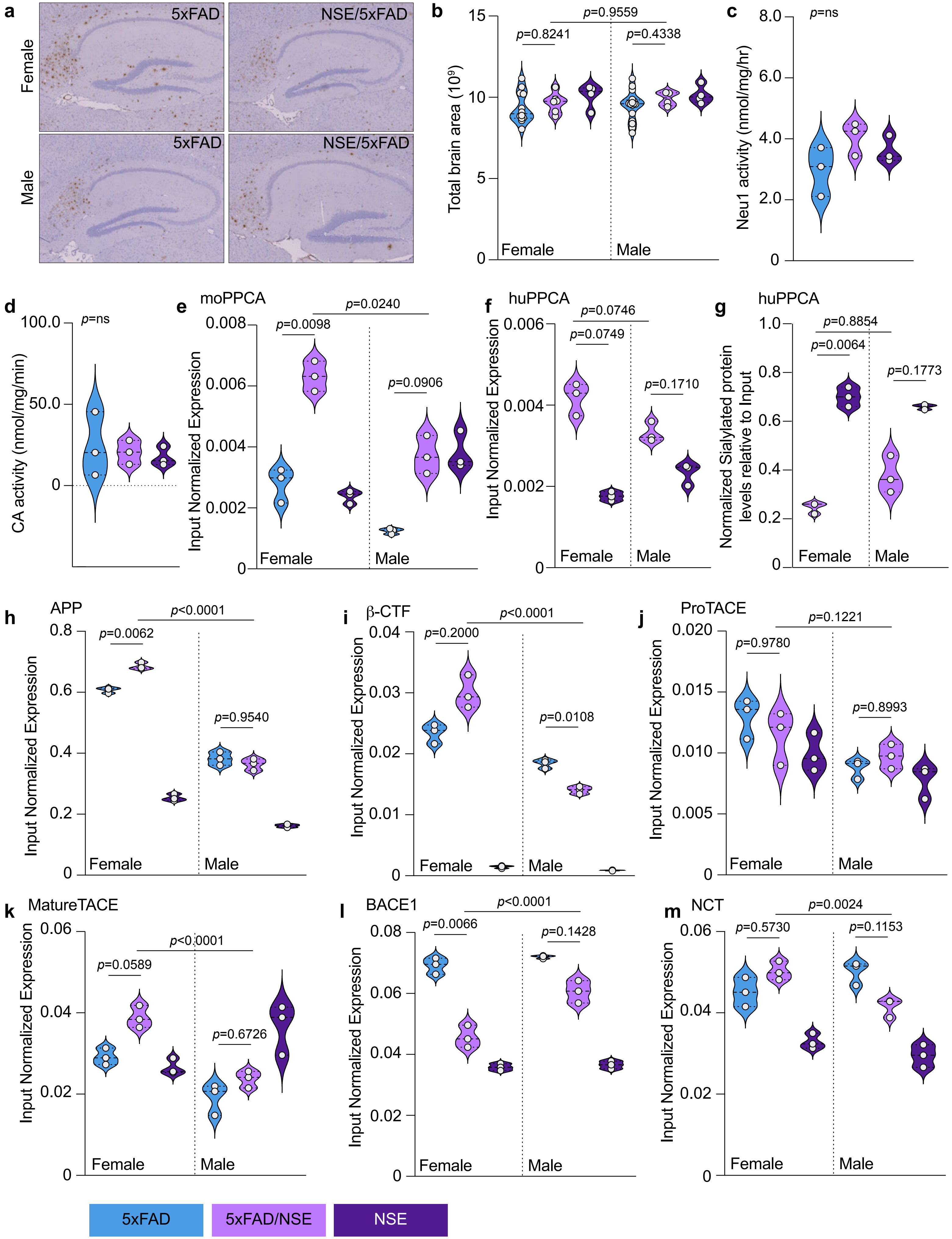
a
Female
Male
5xFAD
NSE/5xFAD
5xFAD
NSE/5xFAD
b
Total brain area (10⁹)
p=0.9559
p=0.8241
p=0.4338
Female
Male
c
Neu1 activity (nmol/mg/hr)
p=ns
d
CA activity (nmol/mg/min)
p=ns
e
moPPCA
Input Normalized Expression
p=0.0240
p=0.0098
p=0.0906
Female
Male
f
huPPCA
Input Normalized Expression
p=0.0746
p=0.0749
p=0.1710
Female
Male
g
huPPCA
Normalized Sialylated protein levels relative to Input
p=0.8854
p=0.0064
p=0.1773
Female
Male
h
APP
Input Normalized Expression
p<0.0001
p=0.0062
p=0.9540
Female
Male
i
β-CTF
Input Normalized Expression
p<0.0001
p=0.2000
p=0.0108
Female
Male
j
ProTACE
Input Normalized Expression
p=0.1221
p=0.9780
p=0.8993
Female
Male
k
MatureTACE
Input Normalized Expression
p<0.0001
p=0.0589
p=0.6726
Female
Male
l
BACE1
p<0.0001
p=0.0066
p=0.1428
Input Normalized Expression
Female
Male
m
NCT
p=0.0024
p=0.5730
p=0.1153
Input Normalized Expression
Female
Male
5xFAD
5xFAD/NSE
NSE

**Extended Data Fig. 5: Analysis of total proteins in 5xFAD mice when PPCA is overexpressed under the NSE promoter.**

**a** and **b**, Collected from paraffin-embedded whole brain sections for female 5xFAD (n=12), female 5xFAD/NSE (n=6), female NSE (n=4), male 5xFAD (n=16), male 5xFAD/NSE (n=4), and male NSE (n=4) mice. **a**, Immunohistochemical staining of amyloid (4G8; brown). **b**, Quantification of total brain area. **c** and **d**, Measurements collected from homogenized hippocampal lysates of 5xFAD, 5xFAD/NSE, and NSE mice (n=3 for all). **c**, Quantification of Neu1 enzyme activity. **d**, Quantification of PPCA (Cathepsin A) enzyme activity. **e-m**, Measurements collected from homogenized hippocampal lysates of female 5xFAD (n=3), female 5xFAD/NSE (n=3), female NSE (n=3), male 5xFAD (n=3), male 5xFAD/NSE (n=3), and male NSE (n=3) mice. **e**, Total normalized protein expression of moPPCA. **f**, Normalized expression of sialylated huPPCA, relative to total protein levels displayed in *Extended Data Fig. 5g*. 5xFAD does not express huPPCA so no data was shown for this group. **g**, Total normalized protein expression of huPPCA. 5xFAD does not express huPPCA so no data was shown for this group. **h-m**, Total normalized protein expression **h**, APP. **i**, β-CTF. **j**, pro-TACE, **k**, mature TACE. **l**, BACE1. **m**, NCT. Actual p-values are given, any value less than or equal to 0.05 is considered significant.

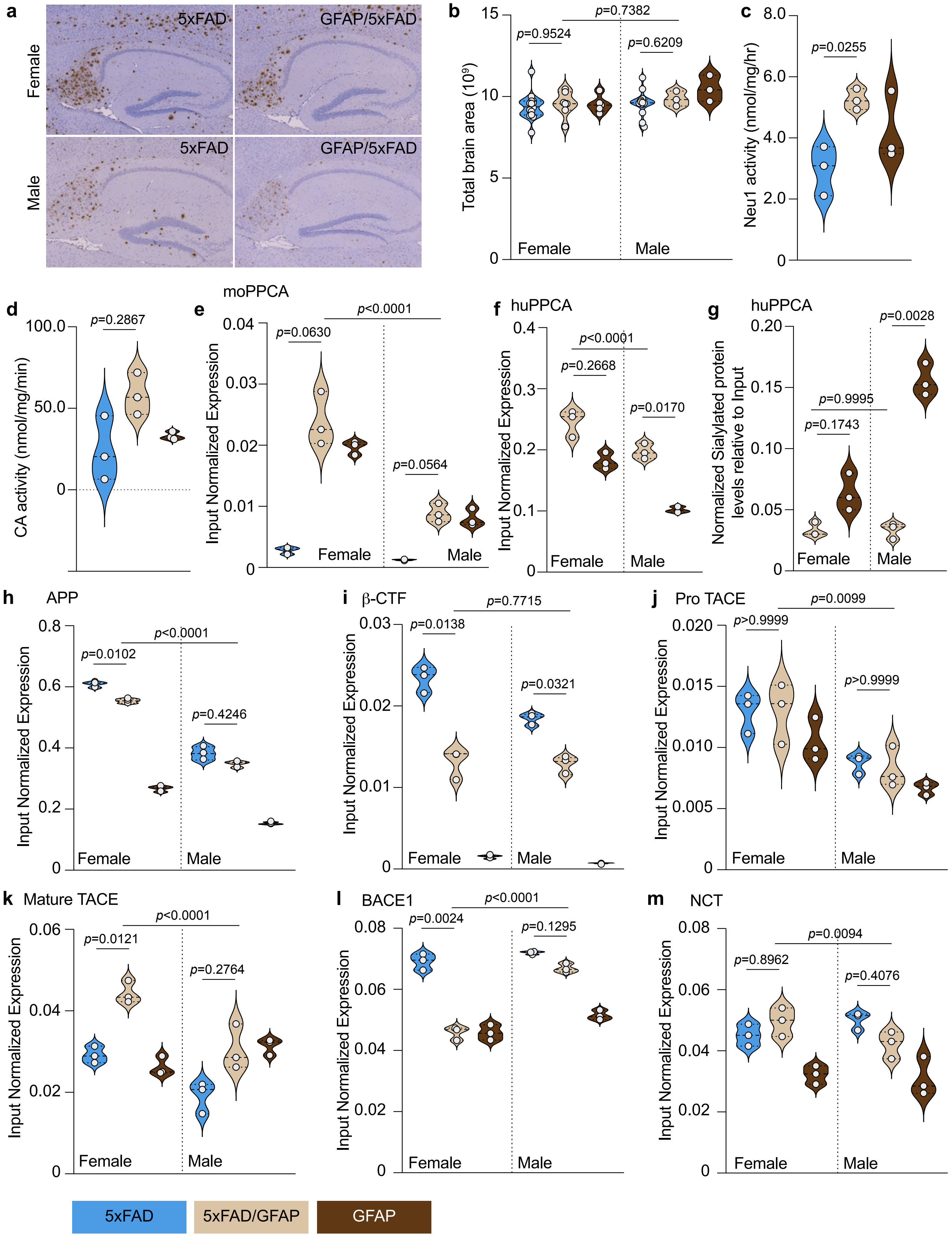
a
Female
Male
5xFAD
GFAP/5xFAD
b
Total brain area ($10^9$)
p=0.9524
p=0.7382
p=0.6209
Female
Male
c
Neu1 activity (nmol/mg/hr)
p=0.0255
d
CA activity (nmol/mg/min)
p=0.2867
e
moPPCA
Input Normalized Expression
p=0.0630
p<0.0001
p=0.0564
Female
Male
f
huPPCA
p<0.0001
p=0.2668
p=0.0170
g
huPPCA
Normalized Sialylated protein levels relative to Input
p=0.0028
p=0.9995
p=0.1743
h
APP
p=0.0102
p<0.0001
p=0.4246
i
β-CTF
p=0.7715
p=0.0138
p=0.0321
j
Pro TACE
p=0.0099
p>0.9999
p>0.9999
k
Mature TACE
p=0.0121
p<0.0001
p=0.2764
l
BACE1
p=0.0024
p<0.0001
p=0.1295
m
NCT
p=0.0094
p=0.8962
p=0.4076
5xFAD
5xFAD/GFAP
GFAP

**Extended Data Fig. 6: Analysis of total proteins in 5xFAD mice when PPCA is overexpressed under the GFAP promoter.**

**a** and **b**, Collected from paraffin-embedded whole brain sections for female 5xFAD (n=14), female 5xFAD/GFAP (n=7), female GFAP (n=5), male 5xFAD (n=12), male 5xFAD/GFAP (n=3), and male GFAP (n=3) mice. **a**, Immunohistochemical staining of amyloid (4G8; brown). **b**, Quantification of total brain area. **c** and **d**, Measurements collected from homogenized hippocampal lysates of 5xFAD, 5xFAD/GFAP, and GFAP mice (n=3 for all). **c**, Quantification of Neu1 enzyme activity. **d**, Quantification of PPCA (Cathepsin A) enzyme activity. **e-m**, Measurements collected from homogenized hippocampal lysates of female 5xFAD (n=3), female 5xFAD/GFAP (n=3), female GFAP (n=3), male 5xFAD (n=3), male 5xFAD/GFAP (n=3), and male GFAP (n=3) mice. **e**, Total normalized protein expression of moPPCA. **f**, Normalized expression of sialylated huPPCA, relative to total protein levels displayed in *Extended Data Fig. 6g*. 5xFAD does not express huPPCA so no data was shown for this group. **g**, Total normalized protein expression of huPPCA. 5xFAD does not express huPPCA so no data was shown for this group. **h-m**, Total normalized protein expression **h**, APP. **i**, β-CTF. **j**, pro-TACE, **k**, mature TACE. **l**, BACE1. **m**, NCT. Actual p-values are given, any value less than or equal to 0.05 is considered significant.

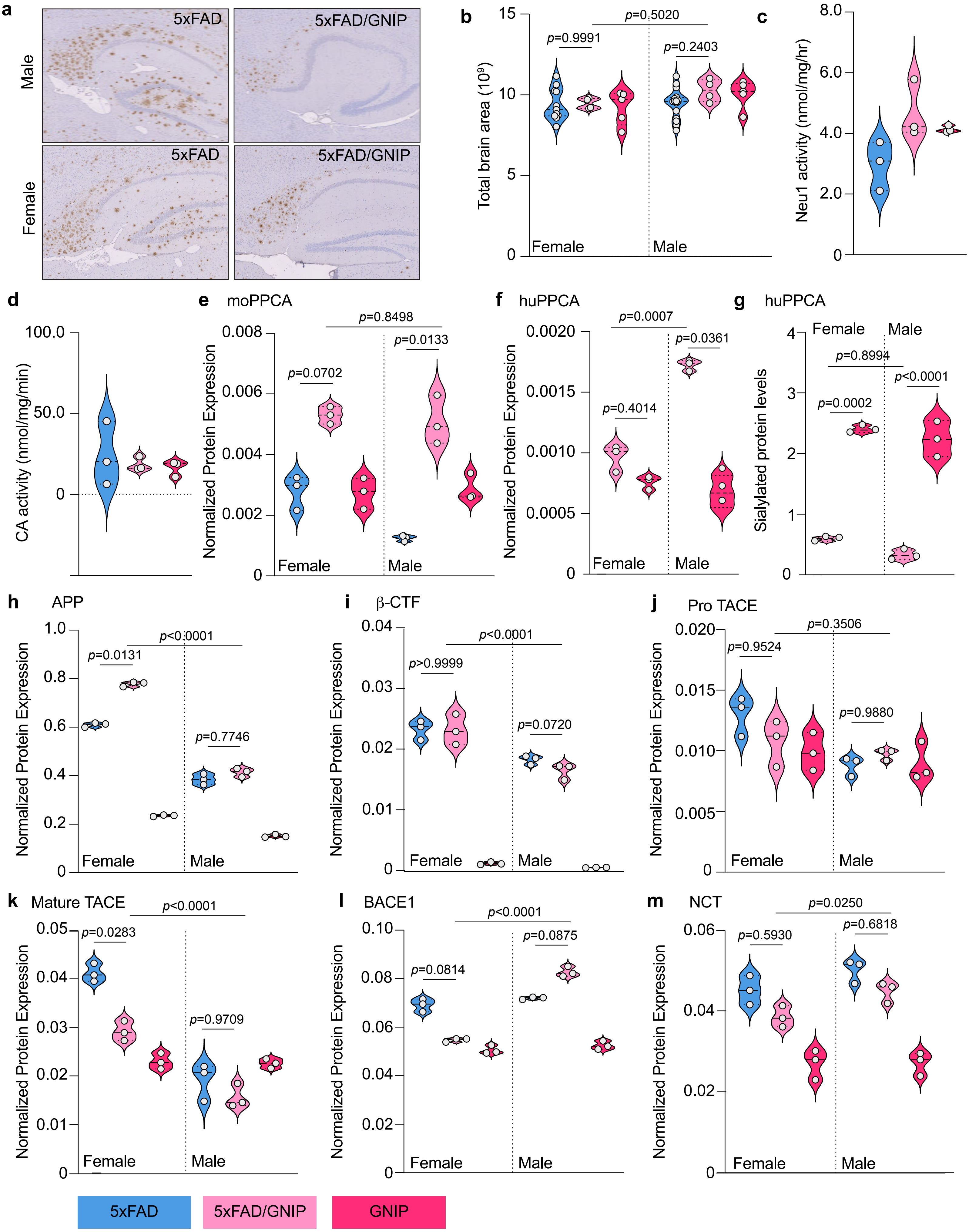

a
Male
Female
5xFAD
5xFAD/GNIP
b
Total brain area (10⁹)
p=0.9991
p=0.5020
p=0.2403
Female
Male
c
Neu1 activity (nmol/mg/hr)
d
CA activity (nmol/mg/min)
e moPPCA
p=0.0702
p=0.8498
p=0.0133
f huPPCA
p=0.4014
p=0.0007
p=0.0361
g huPPCA
Sialylated protein levels
p=0.8994
p=0.0002
p<0.0001
h APP
p=0.0131
p<0.0001
p=0.7746
i β-CTF
p>0.9999
p<0.0001
p=0.0720
j Pro TACE
p=0.9524
p=0.3506
p=0.9880
k Mature TACE
p=0.0283
p<0.0001
p=0.9709
l BACE1
p=0.0814
p<0.0001
p=0.0875
m NCT
p=0.5930
p=0.0250
p=0.6818
Normalized Protein Expression
5xFAD
5xFAD/GNIP
GNIP

**Extended Data Fig. 7: Analysis of total proteins in 5xFAD mice when PPCA and NEU1 are overexpressed under the GFAP promoter.**

**a** and **b**, Collected from paraffin-embedded whole brain sections for female 5xFAD (n=11), female 5xFAD/GNIP (n=4), female GNIP (n=5), male 5xFAD (n=16), male 5xFAD/GNIP (n=4), and male GNIP (n=5) mice. **a**, Immunohistochemical staining of amyloid (4G8; brown). **b**, Quantification of total brain area. **c** and **d**, Measurements collected from homogenized hippocampal lysates of 5xFAD, 5xFAD/GNIP, and GNIP mice (n=3 for all). **c**, Quantification of Neu1 enzyme activity. **d**, Quantification of PPCA (Cathepsin A) enzyme activity. **e-m**, Measurements collected from homogenized hippocampal lysates of female 5xFAD (n=3), female 5xFAD/GNIP (n=3), female GNIP (n=3), male 5xFAD (n=3), male 5xFAD/GNIP (n=3), and male GFAP (n=3) mice. **e**, Total normalized protein expression of moPPCA. **f**, Normalized expression of sialylated huPPCA, relative to total protein levels displayed in *Extended Data Fig. 7g*. 5xFAD does not express huPPCA so no data was shown for this group. **g**, Total normalized protein expression of huPPCA. 5xFAD does not express huPPCA so no data was shown for this group. **h-m**, Total normalized protein expression **h**, APP. **i**, β-CTF. **j**, pro-TACE, **k**, mature TACE. **l**, BACE1. **m**, NCT. Actual p-values are given, any value less than or equal to 0.05 is considered significant.

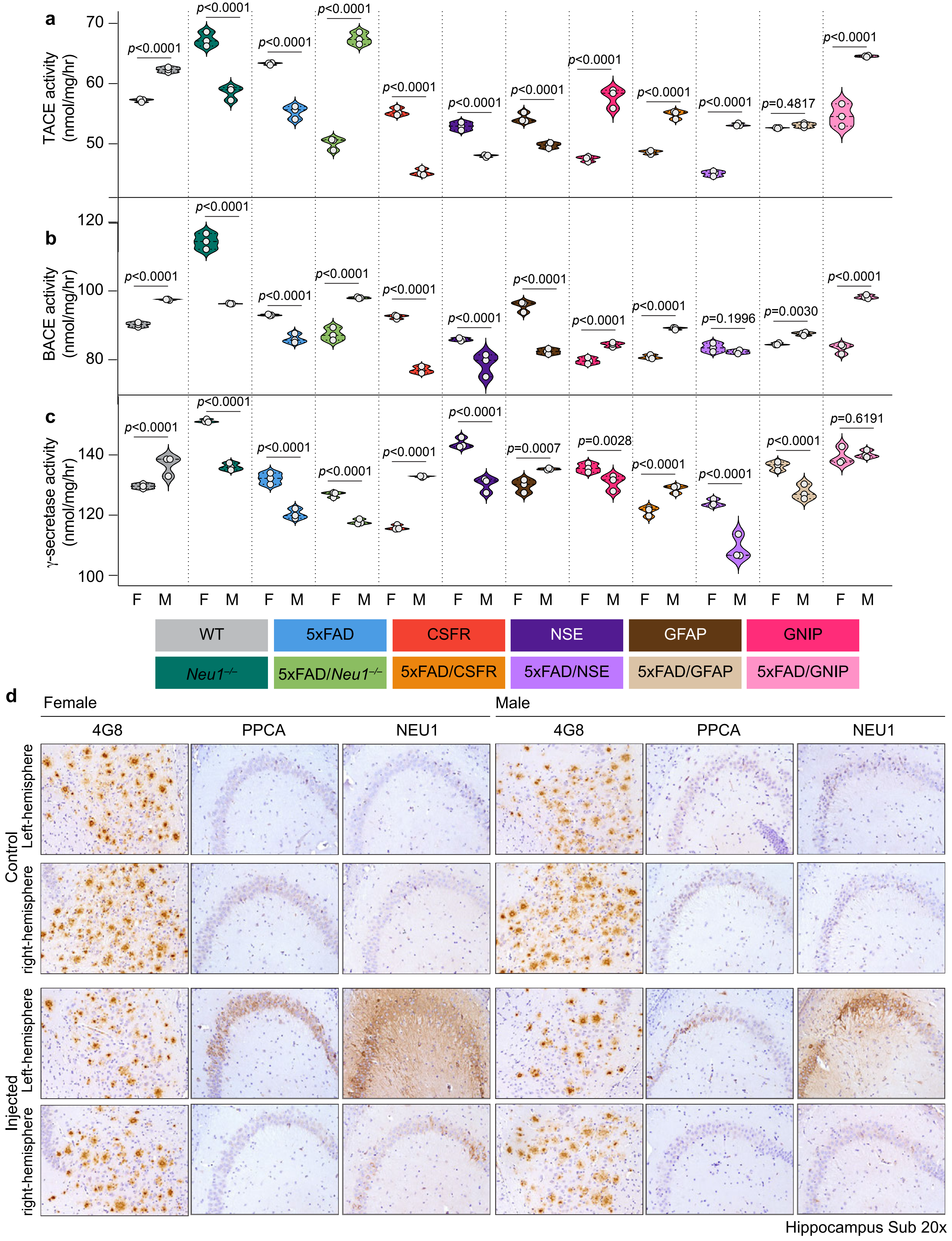
a
TACE activity (nmol/mg/hr)
b
BACE activity (nmol/mg/hr)
c
γ-secretase activity (nmol/mg/hr)
p<0.0001
p=0.4817
p=0.1996
p=0.0030
p=0.0007
p=0.0028
p=0.6191
F M
WT
5xFAD
CSFR
NSE
GFAP
GNIP
Neu1-/-
5xFAD/Neu1-/-
5xFAD/CSFR
5xFAD/NSE
5xFAD/GFAP
5xFAD/GNIP
d
Female
Male
4G8
PPCA
NEU1
Control
Left-hemisphere
right-hemisphere
Injected
Left-hemisphere
right-hemisphere
Hippocampus Sub 20x

**Extended Data Fig. 8: Comparison of female and male secretase activities across the 5xFAD models and visual assessment of AAV-PIN injections in 5xFAD mice.**

**a-c**, Measurements collected from homogenized hippocampal tissue from female and male WT, *Neu1*$^{-/-}$, 5xFAD, 5xFAD/*Neu1*$^{-/-}$, CSFR, NSE, GFAP, GNIP, 5xFAD/CSFR, 5xFAD/NSE, 5xFAD/GFAP, and 5xFAD/GNIP mice (n=3 for all). Female and male plots have been combined for comparison between sexes rather than genetic background. **a**, Quantification of TACE enzyme activity. **b**, Quantification of BACE1 enzyme activity. **c**, Quantification of $\gamma$-secretase enzyme activity. **d**, Images collected from paraffin-embedded whole brain sections for female 5xFAD (n=7), female 5xFAD/AAV-PIN (n=4), male 5xFAD (n=9) and male 5xFAD/AAV-PIN (n=9). Immunohistochemical staining of amyloid (4G8; brown; 1$^{st}$ and 4$^{th}$ panel columns), PPCA (brown; 2$^{nd}$ and 5$^{th}$ panel columns), and NEU1 (brown; 3$^{rd}$ and 6$^{th}$ panel columns) in female (left set of panels) and male (right set of panels) controls (5xFAD; top set of panels) and AAV-injected (5xFAD/AAV-PIN; bottom set of panels) mice. Both left and right hemispheres are shown; AAV injections were made only into left hemispheres, so right hemispheres serve as an internal control.
